# A generic deviance detection principle for cortical On/Off responses, omission response, and mismatch negativity

**DOI:** 10.1101/582437

**Authors:** Shih-Cheng Chien, Burkhard Maess, Thomas R. Knösche

**Keywords:** Deviance detection, Neural mass model, Auditory perception, Adaptation, NMDA

## Abstract

Neural responses to sudden changes can be observed in many parts of the sensory pathways at different organizational levels. For example, deviants that violate regularity at various levels of abstraction can be observed as simple On/Off responses of individual neurons or as cumulative responses of neural populations. The cortical deviance-related responses supporting different functionalities (e.g. gap detection, chunking, etc.) seem unlikely to arise from different function-specific neural circuits, given the relatively uniform and self-similar wiring patterns across cortical areas and spatial scales. Additionally, reciprocal wiring patterns (with heterogeneous combinations of excitatory and inhibitory connections) in the cortex naturally speak in favor of a generic deviance detection principle. Based on this concept, we propose a network model consisting of reciprocally coupled neural masses as a blueprint of a universal change detector. Simulation examples reproduce properties of cortical deviance-related responses including the On/Off responses, the omitted-stimulus response (OSR), and the mismatch negativity (MMN). We propose that the emergence of change detectors relies on the involvement of disinhibition. The analysis on network connection settings further suggests a supportive effect of synaptic adaptation and a destructive effect of N-methyl-D-aspartate receptor (NMDA-r) antagonists on change detection. We conclude that the nature of cortical reciprocal wirings gives rise to a whole range of local change detectors supporting the notion of a generic deviance detection principle. Several testable predictions are provided based on the network model. Notably, we predict that the NMDA-r antagonists would generally dampen the cortical Off response, the cortical OSR, and the MMN.

## 1 Introduction

Automatic detection of sudden acoustic changes crucially enables reorientation of attention towards relevant events in the environment and thereby is important for survival. From a functional perspective, the sensitivity to deviants at the various levels serves different functionalities (e.g. noise rejection, duration tuning, chunking and grouping, beat perception, see reviews in [58,106]) and enriches the hierarchical representations of percepts. The ability to detect abrupt temporal changes is thought to be a pervasive property of the sensory systems, given that deviance-related responses have been widely observed from cellular to system levels, across species, sensory modalities, and spanning from the lower levels of the sensory pathway to the cortex. For example, some cells can be sensitive only to the onsets and offsets of stimuli. These On/Off responses have been observed using extracellular recording in the superior paraolivary nucleus (SPON) of rodents [7,19,21,44,47], inferior colliculus (IC) of chinchillas [24], medial geniculate body (MGB) of the guinea pig [26]. Cortical On/Off responses have been observed using different recording and imaging techniques, including single-cell recording in primary auditory cortex (A1) of awake cats [16,69], and anesthetized rats [80], extracellular recording in A1 of awake marmoset monkeys [76], surface micro-electrode array in auditory cortex (AC) of rats [92], multi-unit extracellular recordings across broad range of AC of mice [39], flavoprotein fluorescence imaging [4] and two-photon calcium imaging [4,20] in AC of mice, and MEG in human auditory evoked responses [61]. Generally speaking, these cells can be sensitive to the sudden changes in specific regular features such as the constancy in pitch, loudness, duration, and patterns. The deviants that violate these perceptual regularities trigger mismatch response at different stages such as frequency following responses (FFR), middle latency responses (MLR), as well as long latency responses (LLR) such as the mismatch negativity (MMN) [85]. An omitted stimulus in a periodic train of stimuli is a special case of deviants, which elicits the so-called omitted-stimulus responses /potentials (OSRs/OSPs). The OSR is time-locked not to the last but to the omitted stimulus, which reflects temporal expectancy represented in the neural circuits. OSRs have been observed in different sensory systems (e.g., visual, auditory, somatosensory) in various species, for example, the visual pathway of fish, reptile, and invertebrate *in vivo* [12,68,71,40], retinas of salamander *in vitro* [83, 104], and the electrosensory system of rays [14]. The OSR at the cortical level (often termed as omission response or omission MMN) has also been observed in human EEG/MEG [29,2,13,41,15]. So far, investigations of the underlying mechanisms have been mostly confined in a certain level and a particular phenomenon. A unifying view of deviance detection that considers phenomena across levels is still missing.

Many of the deviance-related activities, though originating from different stages of the auditory pathway, can be observed pervasively in the auditory cortex. We hypothesize that the cortical deviance-related activities are primarily generated locally through reciprocally connected neural circuits. In this study, we outline a *generic deviance detection* principle, in an effort to reconcile some confusions and conflictions related to the questions below.

### What neural circuits give rise to the diverse cortical On/Off responses?

The response of a neuron or a neural circuit to a sustained stimulus can bear three basic features: a response to the stimulus onset (On response), a sustained response as long the stimulus is present, and a response to the stimulus offset (Off response). The On/Off responses are found in neurons of the superior paraolivary nucleus (SPON) of the brainstem, the inferior colliculus (IC) of the midbrain [22], and the auditory cortex [80,4,20]. These On/Off neurons are thought to support functions such as duration selectivity (duration tuning), gap detection, and noise rejection [106]. Knowledge in the generation of On/Off responses is mainly derived from observations at non-cortical stages. The On responses are thought to be due to adaptive and post-onset inhibitory mechanisms that shape the responses in the auditory nerve [66]. The Off responses are widely accepted to arise from post-inhibitory rebound (see review in [43] for the detailed celluar and synaptic mechanisms), as concluded from observation in SPON neurons [21]. Other response patterns such as On-Off, On-sustained-Off can then potentially be explained by mixing of excitatory and inhibitory inputs with different delays in a feed-forward network [106]. As for the On/Off responses recorded in the auditory cortex, they may originate from the ascending non-cortical On/Off responses [80] or be generated locally in the cortex. The cortical On/Off neurons show diverse temporal profiles [20]. Also, a single cortical neuron may have distinct onset- and offset-frequency receptive fields (FRFs) [69]. It is still unclear how the neural circuits give rise to these properties of cortical On/Off responses.

### Is the OSR just sustained resonance?

The OSR, elicited by an unexpected omission in periodic stimuli, is found in the cortex [29,2,13,41,15], but not in the midbrain (IC, tectum) [61] and the brainstem [50], where only Off responses are observed. The OSR resembles the Off response as they both peak at the end of a stimulus (or a train of stimuli), except that the OSR also reflects temporal expectancy (i.e., neural representation of periodicity), which distinguishes it from the Off response. The properties of the OSR include: (1) The peak latency includes an additional constant delay (e.g., around 100 ms in human MEG/EEG) after the due-time of the missing stimulus which does not depend on the stimulus-onset asynchrony (SOA) of stimuli [2,84]. (2) The peak amplitude can be larger than the entrained responses during periodic stimuli [29]. Althought neural activities that show sustained resonance can be a mechanism underlying the temporal expectancy [52,94], sustained response alone does not explain the additional delay and higher peak amplitude. How the neural circuits maintain the input periodicity and detect the change is unclear.

### Does the OSR reflect prediction or prediction error?

This question rests on whether the OSR is triggered by a similar mechanism as the MMN. The MMN, elicited by a deviant among repetitive standard stimuli, is a negative deflection in the event-related potential (ERP) with the sources most prominently localized in the auditory cortex. The underlying process leads to the reorientation of attention for higher cognitive processes. Deviants which differed in a variety of sound properties from the standard have been shown to elicit MMNs, for example pitch [78,95,56,93,108,109,33,64,63], intensity [57,73], duration [1,75,59,30,57,103,81,32,31,17,36], SOA [46,97,11], sequence (or pattern) [82,28,10,112,45,96], and more complex features such as rising and falling tones (reviewed in [65]) or voice [42]. The MMN is generally accepted to be elicited by the deviant that violates the regularities, but the underlying mechanism is still under debate. The MMN is thought to reflect either the prediction-error signal resulting from the comparison between the input and the top-down prediction (prediction hypothesis), or the fresh stimulus propagating through un-adapted synapses (adaptation hypothesis). The omission paradigms that elicit the OSR are often used in the debate to emphasize the need for active prediction, since the adaptation mechanism alone does not produce extra neural activities without any input. However, according to the computational models based on either hypothesis, the OSR is qualitatively different from the classical MMNs elicited by other deviants. The adaptation-based model suggests the OSR to be a rebound response (i.e. sustained resonance) rather than a modulated N1 [52]. The prediction-based model suggests the OSR to reflect predictive signals rather than prediction error [102]. Both interpretations implicitly suggest pure endogenous activities where a change detection mechanism is not involved, which conflicts with the two properties of ORS mentioned above. The interpretation to MMN generation therefore is not comprehensive yet.

The above issues underscore the need for a unifying view to deviance detection covering the cortical On/Off responses, the cortical OSR, and the MMN. Given the non-specific wiring patterns across areas in the cortex, we ask whether cortical deviance detection can be supported by neural circuits of a common structural motif. We propose a *generic deviance detection* principle (Figure 1a), where change detection can take place locally under proper reciprocal connections (Figure 1b), monitoring the neighboring neural activities that represent a regular feature. This principle is based on the assumption that the process of deviance detection can be functionally separated into stages of *regularity formation* and *change detection.*

**Fig. 1.**
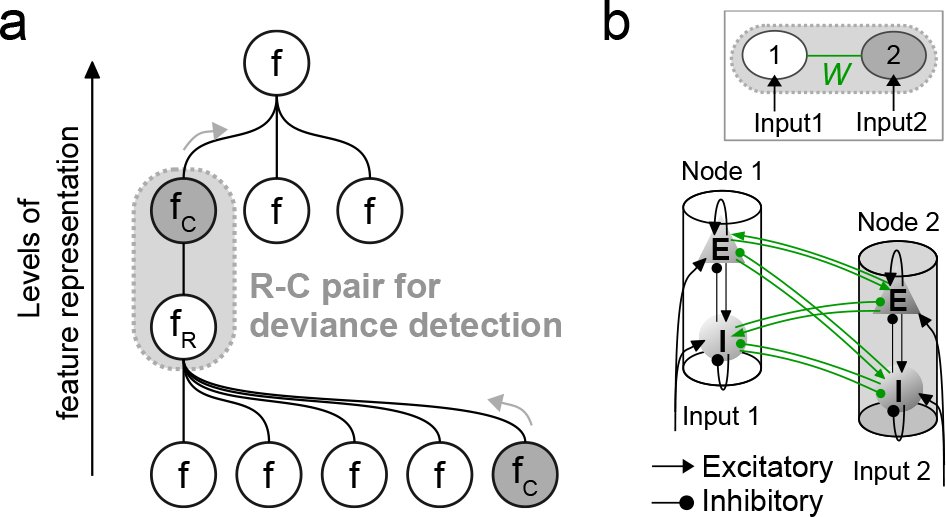
Illustration of the role of deviance detection in hierarchical feature representation. **a** The process of feature representation includes the interaction between regularity formation (R) and change detection (C). The R nodes represent temporally stable feature (regularity, *f*_*R*_) by accumulating and extracting the ascending information from the lower-level features. The C nodes detect abrupt temporal changes in the neighboring R node(s) and pass them as new features (*f*_*C*_, gray arrows) to the higher levels. In this sense, an R-C pair forms a basic mechanism of deviance detection which takes place at every level in the hierarchy. **b** An R-C pair is formed by two reciprocally coupled nodes. In the simulations, all nodes are allowed to receive external weighted inputs that reach the excitatory and inhibitory populations. The internode connections (green) are the free parameters, and the intra-node connections are fixed for simplicity.

In the first part of the Results section, we provide simulation examples that reproduce several properties of cortical On/Off responses, cortical OSR, and MMN. In examples I and II, we demonstrate that the various types of cortical On/Off responses in terms of their temporal profiles and frequency receptive fields (RFRs) can be attributed to the connection patterns between input and observation points. In example III, we demonstrate that the OSR can be regarded as a change detection response (or an Off response) to the cessation of constant periodicity, In example IV, we demonstrate that the sequence MMN can be regarded as change detection responses to the switch in sequence regularity (or a mixture of an On response to the deviant and an Off response to the cessation of regularities). In the second part of the Results section, we study the underlying mechanism of change detection by investigating the generation of simulated On and Off responses. We then study how altered connection patterns (e.g., reduced external connections to inhibitory populations, effect of NMDA-r antagonists, and synaptic adaptation) affect the emergence of change detectors. In the Discussion section, we derive conclusions with regard to the above mentioned questions. Finally, we provide testable predictions for future verification.

## 2 Methods

### 2.1 Model description

The simulations are done with rate-based models which allow for a simple and scalable network motif while keeping the network dynamics comparable to the experimental observations such as LFP, MEG/EEG. In the simulations, a network is used to represent an area in the auditory cortex, and each node in the network comprises one excitatory (*E*) and one inhibitory (*I*) neural population. The dynamics of the *E* and *I* populations are represented by the overall postsynaptic membrane potential (PSP) *υ*^*p*^(*t*) and the mean firing rate *m*^*p*^(*t*), *p* ∈ {*E*, *I*}. Neural populations interact with each other by means of firing rate via connections defined in the matrices *W*^*EE*^, *W*^*IE*^, *W*^*EI*^ and *W*^*II*^ which correspond to excitatory to excitatory, excitatory to inhibitory, inhibitory to excitatory, and inhibitory to inhibitory connections, respectively. Self-feedback is allowed. All *E* populations in the network are fed with constant background input. External stimuli *X*(*t*) reach the *E* and *I* populations via external connections specified by *W*^*EX*^ and *W*^*IX*^.

#### Neural populations

The processing of neural activities in a population is governed by two operators [34,35,88,89]. The *rate-to-potential operator* describes a linear transformation from the mean firing rate to the mean PSP. The input firing rate *x*_*c*_(*t*), *c* ∈ {*e*, *i*} reaches a population via excitatory/inhibitory synapses and is transformed to the EPSP/IPSP *υ*_*c*_(*t*) in that population. This transformation is described by the rate-to-potential process which is achieved by convolving the input firing rate *x*_*c*_(*t*) with a synaptic kernel *h*_*c*_(*t*).

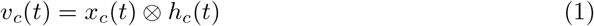

The synaptic kernels *h*_*c*_(*t*) describe the synaptic spike responses and are specified according to different characteristics of the excitatory and inhibitory synapses. In Equation 2, the average synaptic gain *H*_*c*_ controls the maximum of the synaptic spike response curve, and the time constant *τ*_*c*_ represents the delay due to dendritic effects and neurotransmitter kinetics. *Θ*(*t*) denotes the Heaviside step function, where *Θ*(0) = 1.

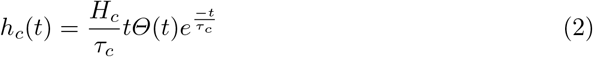

The convolution in Equation 1 can be further represented by two first-order ordinary differential equations:

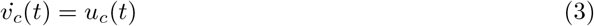

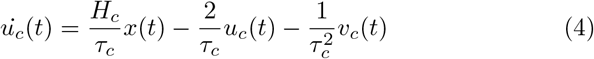

Finally, the overall PSP *υ*(*t*) in the neural population is the summed effect of EPSP and IPSP.

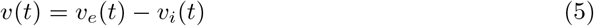

The *potential-to-rate operator* transforms *υ*(*t*) into output firing rate *m*(*t*) by a non-linear sigmoid function *S* as described in Equation 6, where *e*_0_ controls the maximum firing rate and *r* controls the slope at the membrane potential *υ*_0_ for firing.

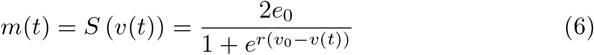

#### Nodes

A node, consisting of one excitatory and one inhibitory neural population, represents the basic building block in a hierarchical feature representation (Figure 1). It represents more as a functional unit rather than a structural one such as a ‘cortical column’. For *N* nodes that represent *N* locations in the auditory cortex and *M* external inputs that represent the intensity of a certain feature such as *M* tones, the network structure is defined by four *N* × *N* connection matrices *W*^*EE*^, *W*^*IE*^, *W*^*EI*^ and *W*^*II*^, and two *N* × *M* external connection matrices *W*^*EX*^ and *W*^*IX*^. Each element *w* (non-negative) in the connection matrices stands for the gain factor between the outgoing and incoming firing rates of the source and target population, respectively. It depends on number and strengths of synapses established between the two populations. The overall PSP of the excitatory population in node 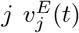 is calculated in Equation 7 as the joint effect of the EPSP 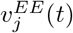 and the IPSP 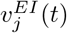

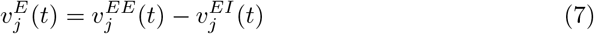

The 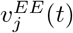 and 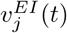 are found by solving the differential equations:

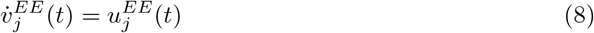

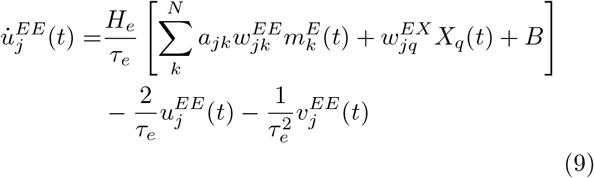

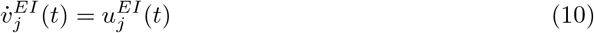

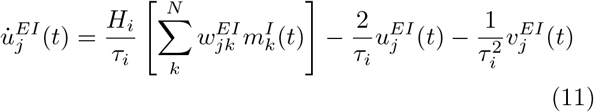

In Equation 9 and 11, 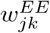, 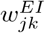 and 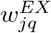 are elements in *W*^*EE*^, *W*^*E1*^ and *W*^*EX*^. The element 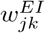 stands for the connection strength from the inhibitory population in node *k* to the excitatory population in node *j*. 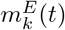 and 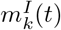 are the outgoing firing rates of the excitatory and inhibitory population in node *k*. *X*_*q*_(*t*) is external input *q*, and *B* is a constant background input. The synaptic adaptation term *a*_*jk*_ modulates the connections strength 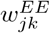. Similarly, the overall PSP of the inhibitory population in node 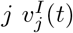 is calculated in Equation 12 as the joint effect of the EPSP 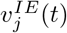 and the IPSP 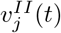.

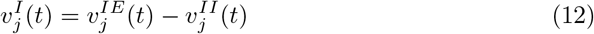

The 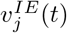 and 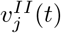 are found by solving the differential equations:

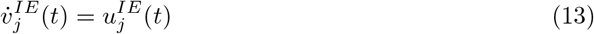

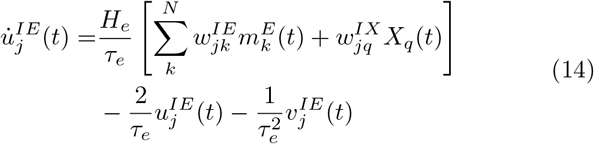

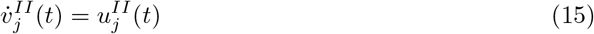

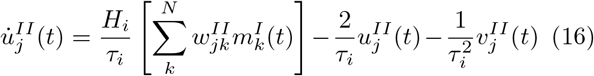

The value of constant background input *B* is chosen so that the nodes work in proper conditions (i.e., near bifurcation point for an isolated node). The external input *X*_*q*_(*t*) reaches both the excitatory and inhibitory populations in node *j* with connection strengths 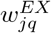 and 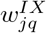, where the ratio 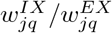 is set to 0.5 by default. The synaptic adaptation term *a* represents the efficacy of excitatory to excitatory connections *W^EE^*. The synaptic efficacy (in range [0,1]) varies according to Equation 17 when synaptic adaptation is considered, otherwise *a* is fixed to 1.

#### Synaptic adaptation

When synaptic adaptation on *W*^*EE*^ is considered, the connection strength 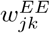 is modulated by the term *a*_*jk*_ in Equation 9 that varies according to the pre-synaptic activity 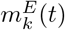.

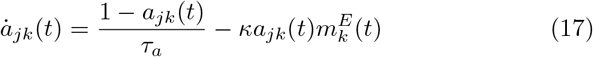

The adaptation time constant *τ*_*a*_ represents the recovery rate of the synaptic efficacy, and the constant *κ* influences the falling speed and the minimum value of *a*_*jk*_(*t*) according to 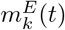.

#### Short-term plasticity

The short-term plasticity is not a focus in this study and is used only in simulation example III as a possible solution for the regularity formation in input periodicity. The plasticity adjusts the bindings among the nodes with different resonance so that the group response maintains a stable representation of input periodicity. The plasticity is applied on 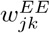 and 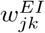, where nodes *j* and *k* are used to encode periodicity. The connection 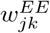 increases if 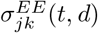, the covariance between 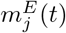 and 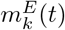 during time *t* − *d* to *t*, is positive and decreases gradually back to zero otherwise (Equations 18,19). Similarly, the connection 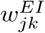 increases if 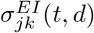 is negative and decreases gradually back to zero otherwise (Equations 20,21). In Equations 19 and 21, *η* is the learning rate. The weights *α*_*jk*_ and *β*_*jk*_ are calculated from two Gaussian functions based on the distance between nodes *j* and *k* (The nodes with similar resonance frequencies are closer to each other). This short-term plasticity learning rule is rather function-driven than based on biological evidence. More studies need to be done for a more realistic network model that maintains the input periodicity.

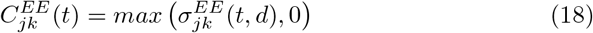

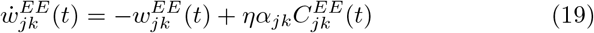

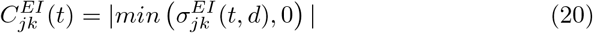

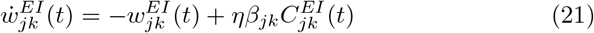

#### Simulated MEG signals

To synthesize a gross signal from the activities of all neural populations in the network, both the excitatory current (or active sink) and inhibitory current (or active source) at the excitatory populations (i.e. Pyramidal cells) are taken into account [87]. This is a more generalized approach than just considering the sum of the excitatory inputs weighted by excitatory to excitatory connection strength and the adaptation term [54]. For the network of *N* nodes, the simulate MEG signal *R*(*t*) is calculated as the weighted sum of currents contributed by the active sinks and sources, in an assumption that the active sinks are due to the EPSP at apical dendrites through *W*^*EE*^, and the active sources are due to be the IPSP at the soma through *W*^*EI*^. In order to highlight the activities of specific nodes (e.g. the change detectors), the signals are weighted by *b*, where 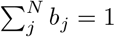.

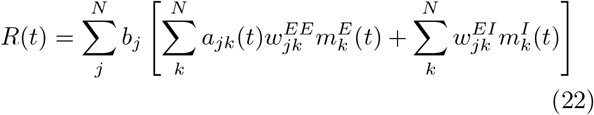

### 2.2 Model configurations

The parameter settings of neural population model are kept the same as proposed by Jansen and Rit model [34,35], unless otherwise specified. In order to reduce the number of free parameters, we fix the intra-node connections and only analyze the inter-node connections in the simulations. The values of intra-node connections are chosen such that a single node stays inactivated under weak excitatory input and starts to oscillate as the excitatory input strength increases to *e*_0_ (i.e., half of the maximum value of the sigmoid function). The adaptation parameters *τ*_*a*_ and *κ* are chosen such that a single node remains oscillating during prolonged stimulation, rather than showing only a transient peak response at the onset. The general configurations are listed in Table 1.

**Table 1.** General configurations

| Par. | Value | Unit | Description |
| --- | --- | --- | --- |
| Impulse response function |  |  |  |
| $\tau_e$ | 10 | ms | Time constant of average delay through excitatory synapses |
| $\tau_i$ | 20 | ms | Time constant of average delay through inhibitory synapses |
| $H_e$ | 3.25 | mV | Average gain through excitatory synapses |
| $H_i$ | 22 | mV | Average gain through inhibitory synapses |
| Sigmoid function |  |  |  |
| $e_0$ | 2.5 | spikes/s | Controlling maximum firing rate |
| $r$ | 0.56 | 1/mV | Slop at $v_0$ |
| $v_0$ | 6 | mV | Membrane potential threshold for firing |
| Intra-node connections |  |  |  |
| $w_{jj}^{EE}$ | $135 \times 0.8$ | 1 | Self-feedback excitatory synapses in the $j$th excitatory population |
| $w_{jj}^{IE}$ | $135 \times 0.6$ | 1 | Excitatory synapses from the $j$th excitatory to the $j$th inhibitory population |
| $w_{jj}^{EI}$ | $135 \times 0.2$ | 1 | Inhibitory synapses from the $j$th inhibitory to the $j$th excitatory population |
| $w_{jj}^{II}$ | $135 \times 0.05$ | 1 | Self-feedback inhibitory synapses in the $j$th inhibitory population |
| Inter-node connections |  |  |  |
| $w_{jk}^{EE}$ | $135 \times \{0, 0.1, \dots, 0.5\}$ | 1 | Excitatory synapses from the $k$th excitatory population to the $j$th excitatory population |
| $w_{jk}^{IE}$ | $135 \times \{0, 0.1, \dots, 0.5\}$ | 1 | Excitatory synapses from the $k$th excitatory population to the $j$th inhibitory population |
| $w_{jk}^{EI}$ | $135 \times \{0, 0.1, 0.2\}$ | 1 | Inhibitory synapses from the $k$th inhibitory population to the $j$th excitatory population |
| $w_{jk}^{II}$ | $135 \times \{0, 0.1, 0.2\}$ | 1 | Inhibitory synapses from the $k$th inhibitory population to the $j$th inhibitory population |
| External connections |  |  |  |
| $w_{jq}^{EX}$ | $220 \times 0.2$ | 1 | Excitatory synapses from external input $q$ to the $j$th excitatory population |
| $w_{jq}^{IX}$ | $220 \times 0.1$ | 1 | Excitatory synapses from external input $q$ to the $j$th inhibitory population |
| Inputs |  |  |  |
| $B$ | $220 \times 2.5$ | spikes/s | Constant background input |
| $X_q(t)$ | - | spikes/s | External input $q$ (step function, amplitude=1.5, rise/fall time=10ms) |
| Synaptic adaptation |  |  |  |
| $\tau_a$ | 200 | ms | Time constant of recovery rate of synaptic efficacy |
| $\kappa$ | 2 | 1 | Drop rate in synaptic efficacy |

### 2.3 Categorization of network behavior

In a two-node network where a prolonged stimulus (2000 ms) is fed to node 1 (Figure 2a), the behavior of node 2 (i.e., the time course 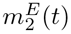) is categorized into one of the eight types based on the level changes and the peak at edges: (1) sustained-Pure, (2) sustained-On, (3) sustained-Off, (4) sustained-On&Off, (5) inhibited-Pure, (6) inhibited-On, (7) inhibited-Off, and (8) inhibited-On&Off. The ‘sustained’ and ‘inhibited’ stand for increased and decreased activity during the stimulus. The ‘On’, ‘Off’ and ‘On&Off’ stand for transient peak(s) at only the onset, only the offset and both the onset&offset of the stimulus. The ‘Pure’ stands for no clear peaks at the edges of the stimulus. Bistable behaviors are not included in the categorization. (See Figure 2b and Table 2 for details of categorization.)

**Fig. 2.**
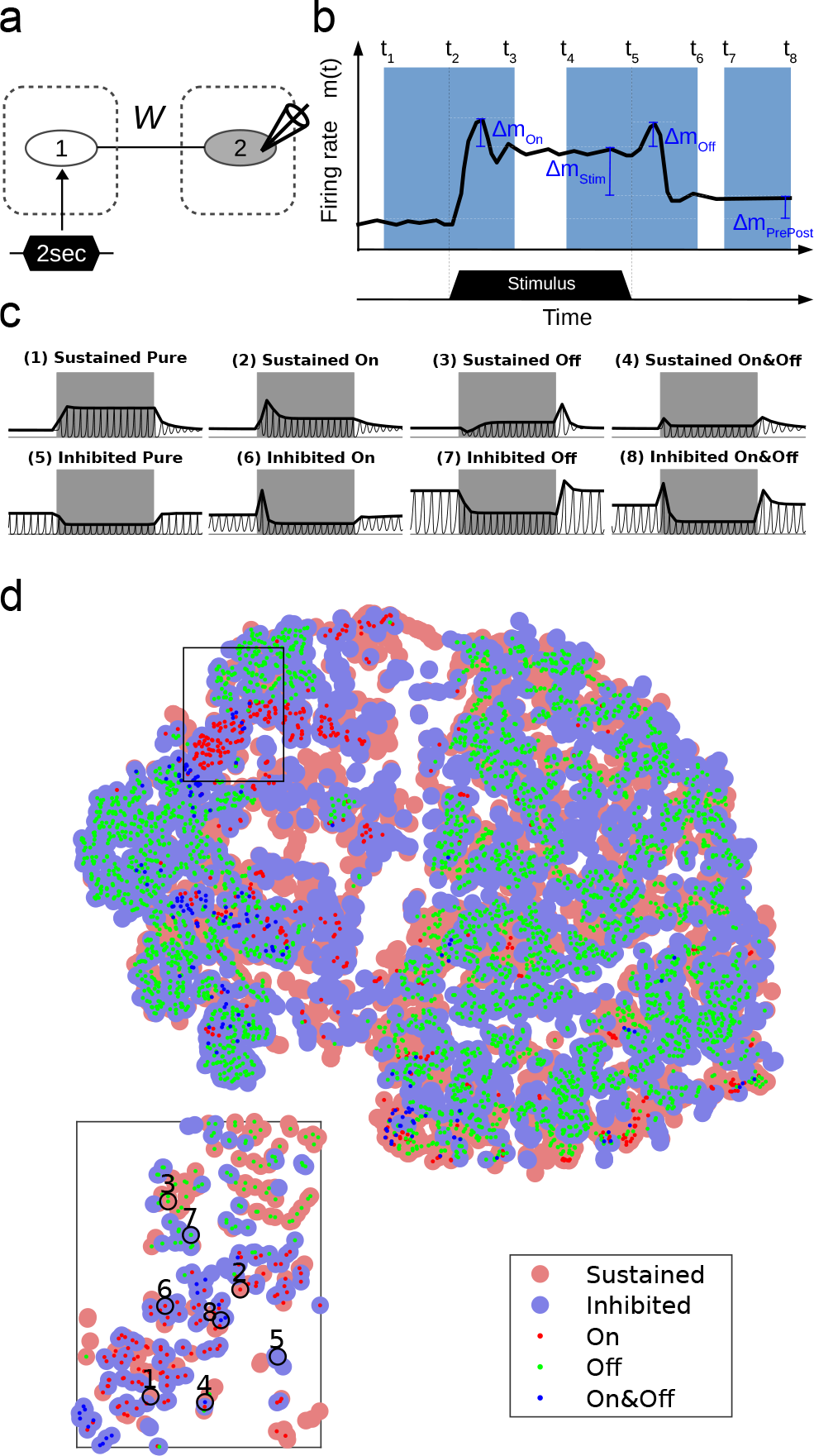
Change detectors and the corresponding *W* solutions. **a** In the simulation settings, a prolonged stimulus of 2000 ms is fed to node 1. A range of inter-node connections *W* are scanned through and the various temporal behaviors of the change detector (i.e., time courses of 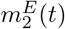) are to be categorized. **b** For categorization, four variables 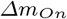, 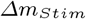, 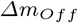 and 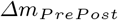 are calculated according to the time windows (light blue areas) for each time course 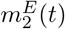. The time courses that are not bistable are then categorized into one of the eight On/Off types. See detailed categorization settings in Table 2. **c** The exemplary responses of eight On/Off types. Gray bands represent the duration of stimulus. The black curves represent the time courses of 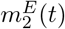, and the bold black curves are the envelopes. **d** All *W* solutions of eight On/Off types in the scanned range are projected to 2D plane for visualization (Matlab function: tsne), where color dots represent the eight types ({Sustained, Inhibited} × {Pure, On, Off, On&Off}). The eight exemplary behaviors in **c** are labeled in the zoomed area.

**Table 2.**
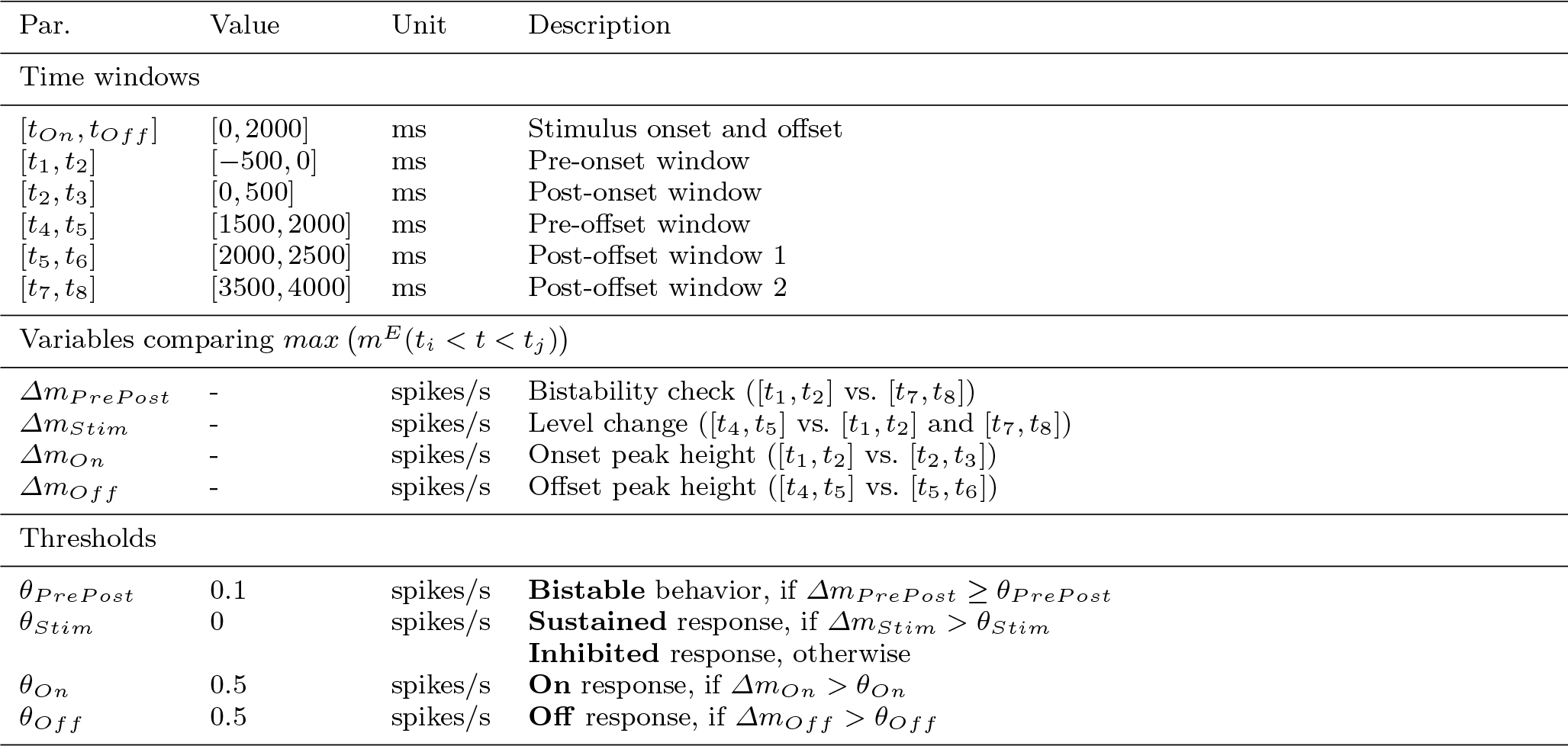
Settings and variables for categorization of network behavior

## 3 Results

The generic deviance detection principle suggests that deviance detections take place locally in the perceptual hierarchy as illustrated in Figure 1. Any two reciprocally coupled nodes in a network can potentially form an R-C pair that serves deviance detection. The connections within an R-C pair can be heterogeneous across locations, thus giving rise to various behaviors of change detectors. Below, we reproduce some observed phenomena of deviance-related responses using simple networks (e.g., comprising two, three, and twenty one nodes) in simulation examples (Section 3.1), and then we investigate the behavior of change detectors and the corresponding network settings in Section 3.2.

### 3.1 Simulation examples

#### Example I: temporal profiles of cortical On/Off responses

A sustained tone stimulus can elicit diverse temporal patterns of On/Off responses in the auditory cortex. Neurons can be sensitive to the onset/offset of the stimulus (i.e., transient responses at the edges), and meanwhile also show increased or decreased firing rate during the stimulus (i.e., level changes) compared with the spontaneous activity [16,69,39,20,72,100]. In this simulation, we feed the input stimulus (2000 ms duration) to a two-node network (Figure 2a), where the change detector does not directly receive the input stimulus (i.e., the external connections to node 2, 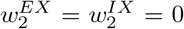). Varying the inter-node connections *W* alters the response of the change detector (e.g., the firing rate of its excitatory population 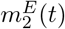). We scan through a range of inter-node connections (*W*^*EE*^, *W*^*IE*^ ∈ {0,0.1,…, 0.5}; *W*^*EI*^, *W*^*II*^ ∈ {0,0.1, 0.2}), and categorize the time courses of 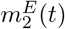 into one of the eight types based on the level changes and the peak at edges (Figure 2b). The *W* solutions are connection settings that give rise to one of the eight categorized On/Off types under this specific simulation settings (e.g., the intensity and onset/offset time of stimulus, the intensity of background input, and intra-node connections, etc).

To further investigate the relation between the internode connections *W* and the On/Off responses, we project the *W* solutions {*W*_*type_i*_,*i* = 1, 2,…, 8} to a 2D plane by t-Distributed Stochastic Neighbor Embedding [51] to visualize the mutual proximity of *W* solutions in the original eight-dimensional space. From Figure 2d we observe that: (1) Although the *W* solutions show clustered pattern from a broader scale, different types are found to be mixed in space when zooming in. The clustering patterns and its sensitivity to *W* may potentially explain the diverse but spatially clustered On/Off responses shown in Figure 5 of [20]. (2) The Off types are not constrained within sustained/inhibited clusters, suggesting that Off responses are not crucially determined by the level change of 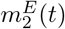 during stimulus. (3) The On and Off types occupy distinct areas in the 2D plane, which agrees with the conclusion that On and Off responses are driven by largely nonoverlapping sets of synaptic inputs [80]. (4) However, there are also areas where the On, Off and On&Off types are close to each other, which supposedly are more sensitive to neuroplasticity (e.g. synaptic adaptation, spike-timing-dependent plasticity, or homeostatic plasticity) that change neural response from one type to another.

#### Example II: distinct onset and offset frequency receptive fields (FRFs)

As demonstrated in Example I, the two-node network can account for the different temporal profiles of On/Off responses. The same network can account for the distinct onset and offset FRFs in individual cells in the auditory cortex [69]. For example, the exemplary cell in Figure 3a is sensitive to the onsets of sound stimuli at higher frequencies (3,200-15,872 Hz) and the offsets of sound stimuli at lower frequencies (512-16,000 Hz), as reflected by higher spike density (yellow and red). In addition, this cell shows suppressed spike density (deep blue) during stimuli at low and middle frequencies. In short, the On/Off responses vary across tonal frequencies and across cells.

**Fig. 3.**
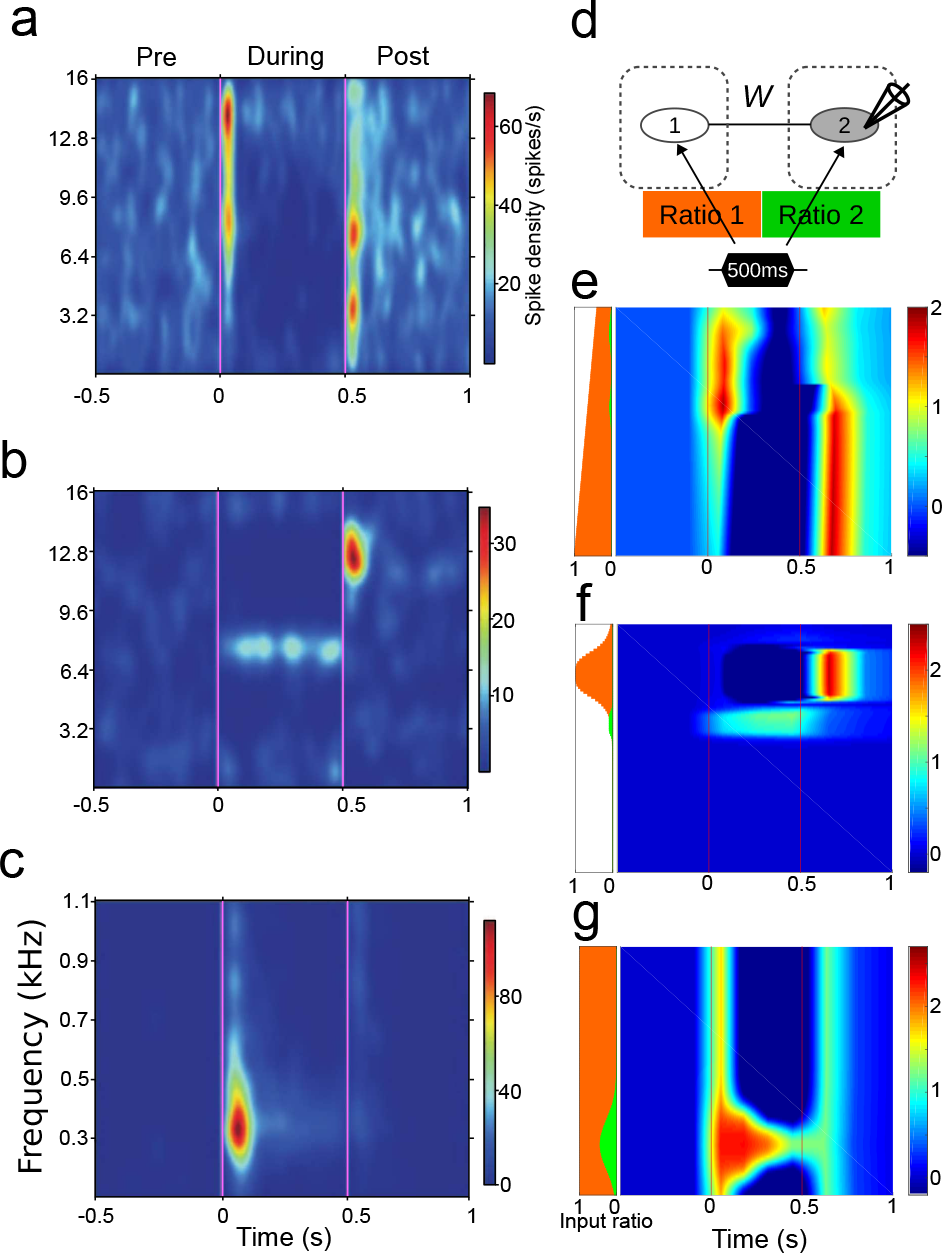
Distinct onset and offset FRFs. **a-c** Three exemplary cells that show distinct onset and offset FRFs (adapted from [69]). The cells were recorded in the primary auditory cortex in awake cats. Sound stimuli of pure tone (ranging from 128Hz to 16,000Hz) were presented for 500 ms. The pre-, during-, and post-stimulus spike density of the cell is color coded. **d** In the simulation settings, a two-node network with adjustable external connections 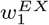 and 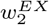 (orange and green color) are used to mimic the experimental observations. **e-g** Simulation results that mimic the observations in **a-c**. The green and orange input ratios at the left-side bar of each plot represent the settings of external connections 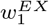 and 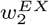 for each simulation trial. The firing rate is color coded. The pre-stimulus firing rate is used as baseline, and the negative value (deep blue color) during stimulus stands for inhibited activity.

In the simulation, we use the two-node network to reproduce the distinct FRFs in Figure 3a-c. For each simulation trial, the stimulus input (500 ms duration), corresponding to a pure tone in one trial of the experimental recordings, is fed to both nodes with different strength of external connections (i.e., 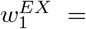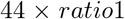; 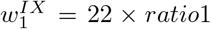; 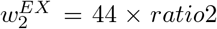; 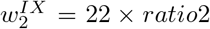, as in 3d). The ratios (orange and green) reflect how far the two nodes are from the stimulus source. Considering the tonotopic organization in the auditory cortex, the ratios are also changed for each simulation trial because the stimulus input in each simulation trial represents a different tonal frequency. The inter-node connections *W* is picked up from the *W* solutions and is fixed in each example, and the ratios are adjusted such that the responses of node 2 (i.e., the time courses of 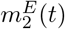 mimic qualitatively the experimental observations. The simulation trials are then merged to make simulated FRFs (Figure 3e-g).

In Figure 3e, the excitatory population *E*_2_ shows inhibited-Off response when *ratio*1 = 1 and *ratio*2 = 0 (same as the ideal case used in Example I). The On response emerges as ratiol decreases, and a small amount of *ratio*2 results in stronger On responses and weaker Off responses. In Figure 3f, *E*_2_ shows inhibited-Off response when *ratio*1 = 1 and *ratio*2 = 0, and it turns into sustained-Pure type when *ratio*2 is larger than *ratio*1. In Figure 3g, *E*_2_ shows inhibited-On&Off response, and a small amount of *ratio*2 results in stronger On responses.

The two-node network, although rate-based, may provide a sense how the exemplary cells in Figure 3a-c are influenced by different sound tones: *ratio*2 (green) indicates which tones are closer to (or more directly influencing) the cell, and *ratio*1 (orange) reflects how its surrounding neurons are sensitive to the tonal scope.

#### Example III: omitted-stimulus response (OSR)

The OSR resembles the Off response as they both peak at the offset of a prolonged stimulus or a train of periodic stimuli. However, the OSR differentiates itself from the Off response by its property of temporal expectation. The peak latencies of OSR are not constant but proportional to the stimulus onset asynchrony (SOA) of the repetitive stimuli as illustrated in Figure 4a. The OSR at cortical level (aka. omission response or omission MMN) resembles the classic MMN, as both responses are related to violations to certain expectations (e.g. expectation to the ‘when’ or the ‘what’ in the stimuli).

**Fig. 4.**
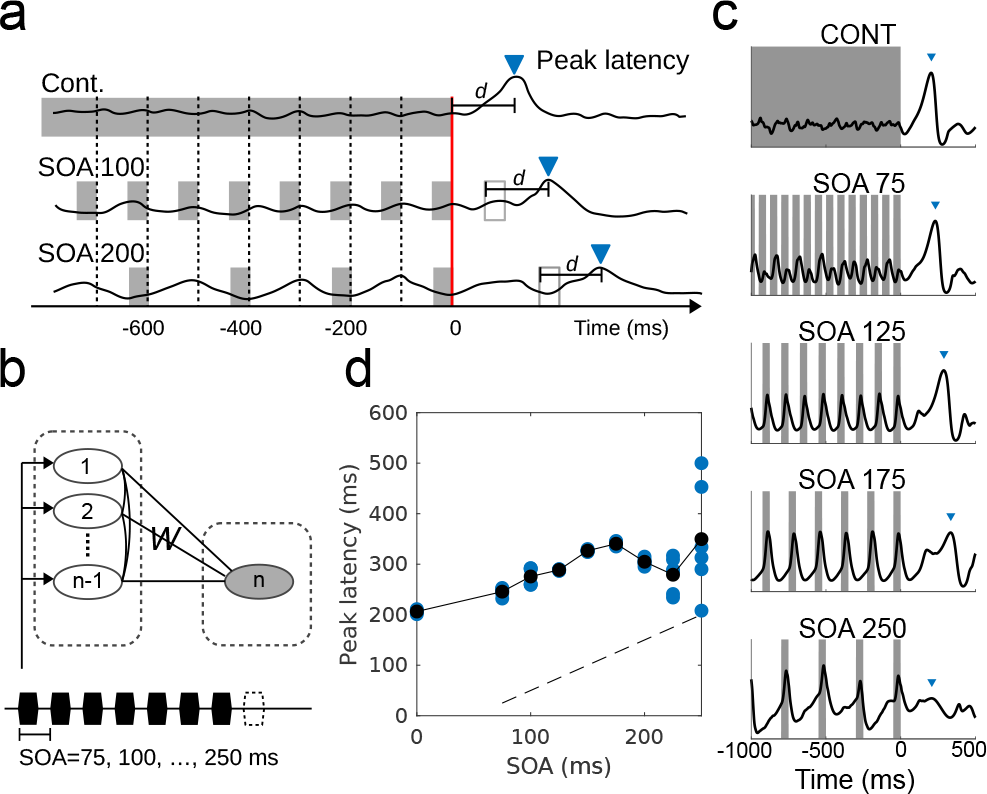
Omitted stimulus response (OSR). **a** Illustrative responses that show temporal expectation. The peak latencies should be linear to the SOA if the offsets of stimuli are aligned (red line), or a constant d if the due times are aligned (empty rectangles). See, for example, Figure 2 in [2]. **b** In the simulation settings, the R nodes (left column) are simply implemented with different time constants *τ*_*e*_ and *τ*_*i*_, leading to different resonance frequencies. A prolonged stimulus or periodic stimuli are fed to these R nodes. **c** Simulated MEG signals (black curves) peak at the end of stimuli and show different peak latencies and peak amplitudes (marked with blue triangles). **d** Simulated peak latencies are linear to the SOA. Simulations are run for several trials for each SOA where the offset time is changed. The peak latencies in each simulation trial (blue dots) under the same SOA can be different, which depends on the network stability during stimulus and the offset time. Black dots are the mean peak latencies. The peak latencies show approximately constant delay with respect to due time (time of predictable omission, dashed line) when the SOAs are below 200 ms. The peak latencies become unstable across trials when SOAs are above 200 ms. In other words, the temporal expectation is preserved in this network for SOAs smaller than 200 ms.

The generic deviance detection principle suggests that the cortical OSR is a change detection response (or an Off response) to the end of stable periodicity representation. In our simulation, the periodicity is represented by a *bank of oscillators* [52,48] comprising multiple nodes (i.e. R nodes) with different resonant frequencies (implemented by different time constants *τ*_*e*_ and *τ*_*i*_ for simplicity). In this way, the temporal feature of periodicity is transformed into a spatial pattern represented by the R nodes. Unlike the conventional bank of oscillators where the oscillators are not connected to each other, the R nodes are inter-connected with short-term plasticity on *W*^*EE*^ and *W*^*E1*^ (Note: short-term plasticity on *W*^*EE*^ and *W*^*IE*^ also works in this example). The plasticity enhances the connections between two nodes if they oscillate in high covariance, and reduces the connections otherwise (Equation 19 and 21). This enables the resonance among R nodes to sustain for one more cycle after the periodic stimuli are turned off. The change detector (C node) that connects to the R nodes (as in Figure 4b) is expected to peak when the sustained resonance drops. In Figure 4c, we simulate MEG signals resulting from prolonged (CONST) and periodic stimuli (SOAs: 75, 125, 175, and 250 ms; stimulus duration: 50 ms). The OSR peaks are marked by blue triangles. When the SOA is increased, the peak latency increases and the peak amplitude decreases, which is in line with the MEG observation [2]. The small peak before the OSR (especially clear for SOA 125 and 175) locates at the time of the omitted stimulus, which resembles the expected evoked potential before the OSR (e.g., Figure 7B in [12]). In Figure 4d, we show that the n-node network (*n* = 21 in this example) is able to respond at the right timing (i.e. a constant delay after the detectable omission) with the limitation that the SOA is within 150 ms. The peak latencies become unstable for SOAs larger than 200 ms. The limitation is due to the limit of resonant frequencies in the bank of oscillators, as can be seen in Figure 4c the simulated MEG for SOA 250 is not as stable compared to faster SOAs.

In this example, we demonstrate that the cortical OSR can reflect a detection mechanism upon the stable representation of periodicity. The sustained resonance is crucial for temporal expectation. This is in line with the observation that the auditory brainstem does not generate overt OSR [50] because sustained resonance has not happened at that stage. We have not yet fully investigated the neural mechanism underlying temporal expectancy. The auditory cortex is assumed to have the capacity to represent a certain range of periodicity locally (e.g. under 200 ms), and the bank of oscillators, which only assumes heterogeneity across neural populations, is so far a good candidate for implementation.

#### Example IV: sequence mismatch negativity (MMN)

The responses in a roving paradigm reveal the progress of regularity formation and change detection, and thus is a good example for demonstrating the generic deviance detection principle. In Figure 5a, an MEG study shows how the human brain responds to the switch between regular and random complex acoustic patterns [5]. There are On and Off responses at the onsets and offsets of the stimulus sequence. An MMN response is elicited by the transition from regular to random sequences (REG-RAND), but there is only gradually rising root mean square (RMS) amplitude for the other way around (RAND-REG). Also, the RMS amplitude is higher during regular sequences compared to random sequences.

**Fig. 5.**
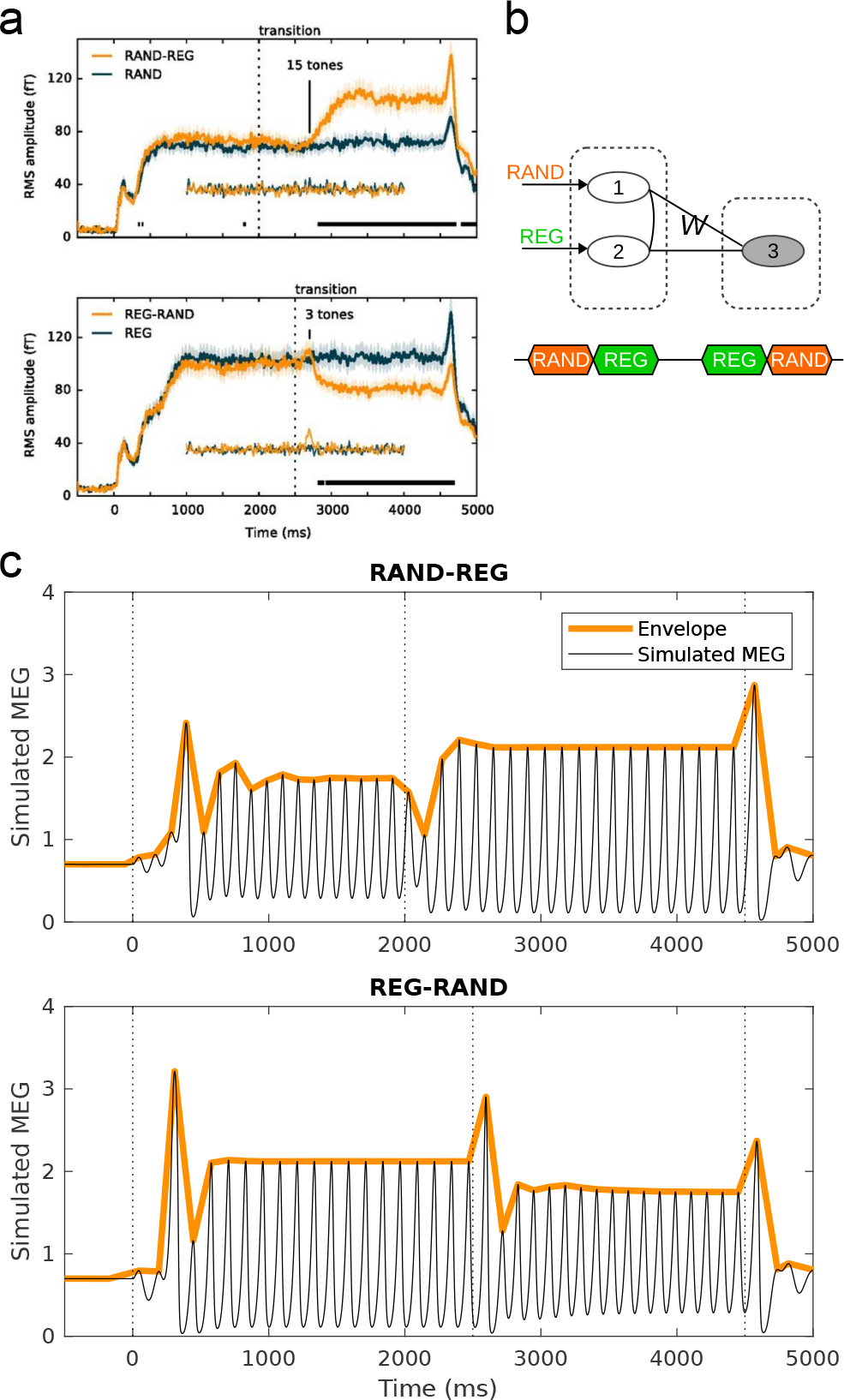
Sequence MMN (roving paradigm). **a** Brain responses to transitions between regular and random sound sequences (adapted from [5]). There are transient peaks at the onsets/offsets of sound sequences as well as at the transition from regular (REG) to random (RAND) sequences. In addition, the RMS amplitude is higher during regular sequences. **b** In the simulation settings, a three-node network is used for mimicking the observation in **a**. The R nodes (nodes 1 and 2) receive the stimulus inputs representing RAND and REG, respectively. The inter-node connections *W* between the C node (node 3) and the R nodes are picked up from the W solutions such that the C node shows sustained-On&Off responses to the stimulus inputs. The connections between the R are tuned to result in the different level shifts during stimulus and the elimination of the transient peak at the transition from RAND to REG. **c** Simulated MEG signals of the three-node network.

In the simulation, we use a three-node network to reproduce the temporal profile of the RMS in Figure 5a. Two stimulus inputs (REG and RAND) that represent the random and regular features are fed to the nodes 1 and 2 respectively as in Figure 5b. The intensity and rise/fall time of the two stimulus inputs are the same as in the previous examples, and the durations are set to match the experiment. The inter-node connections *W* between nodes 1,2 and node 3 are chosen from the *W* solutions in Figure 2d. The connections between nodes 1 and 2 do not have to be symmetric and are manually tuned to match the observed RMS. In Figure 5c, the simulated MEG signal shows (1) On and Off responses at the onset and offset of stimulus sequences. (2) MMN response to the transition from regular to random sequences (REG-RAND), and (3) different RMS amplitudes during REG and RAND presentations.

The three-node network demonstrates how the internode connections *W* among the three nodes alone can account for the transient responses to the onsets and offsets, and the selectivity to the direction of transition, as well as the level changes in RMS amplitude during random or regular sequences. For more realistic settings, the rise/fall time of the two stimulus inputs can be set differently. For example, it is reasonable to set longer rise time for the REG stimulus input because it takes some time (at least a sequence length) to form regularity representation, which then already explains why there is no MMN in the RAND-REG transition. Moreover, the intensity of the two stimulus inputs may be reasonably set differently because the status of neural populations under regular and random sequences can be dramatically different, which then explains the level changes in RMS amplitude. In this simulation example we use identical stimulus inputs, in an attempt to highlight the effect of inter-node connections *W* on shaping the network activities. Note that this simulation example elucidates on the contribution of a change detector, rather than the details of regularity formation. To understand how REG sequence causes higher RMS amplitude, we assume short-term plasticity on *W*^*EE*^ and *W*^*IE*^ in the lower-level neural populations at the stage of regularity formation. This follows the suggestion by a dynamic causal modeling study [3] that synaptic gain modulation in the auditory cortex is involved in processing regular sequences.

### 3.2 The requirements for a change detector

The generic deviance detection principle emphasizes on the ubiquity of local change detection and its separation from regularity formation. In previous simulation examples, we demonstrate that the behavior of a change detector can account for many phenomena (e.g., diverse cortical On/Off responses, distinct onset- and offset-FRFs, cortical OSR, and sequence MMN). Here, we present a more detailed analysis of the exact requirements for a change detector to work. First, we investigate how and under which conditions the On and Off responses occur. Then we examine how changes in connection strengths affects the generation of On/Off responses through three factors: (1) external input to inhibitory populations, (2) blockage of NMDA receptor channels, and (3) synaptic adaptation.

#### The generation of On responses

It has been proposed that On responses can be due to adaptive and post-onset inhibitory mechanisms that reshape the onset response in auditory nerve fibers [66]. In our simulations, we found that the On responses can be also due to the transiently inhibited activity of the inhibitory population *I*_2_ at the onset of stimulus. As can be seen in Figure 6b and d, population *I*_2_ is shortly inhibited by population *I*_1_, and the low 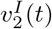 leads to a transient peak in 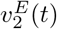 (indicated by the red and black arrows in the magenta rectangles). The system returns to stability soon after the 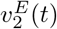 peak brings 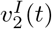 up again. These On responses are due to the *transient disinhibition*, therefore the inter-node connection *W*^*II*^ plays an important role in the generation of the On responses.

**Fig. 6.**
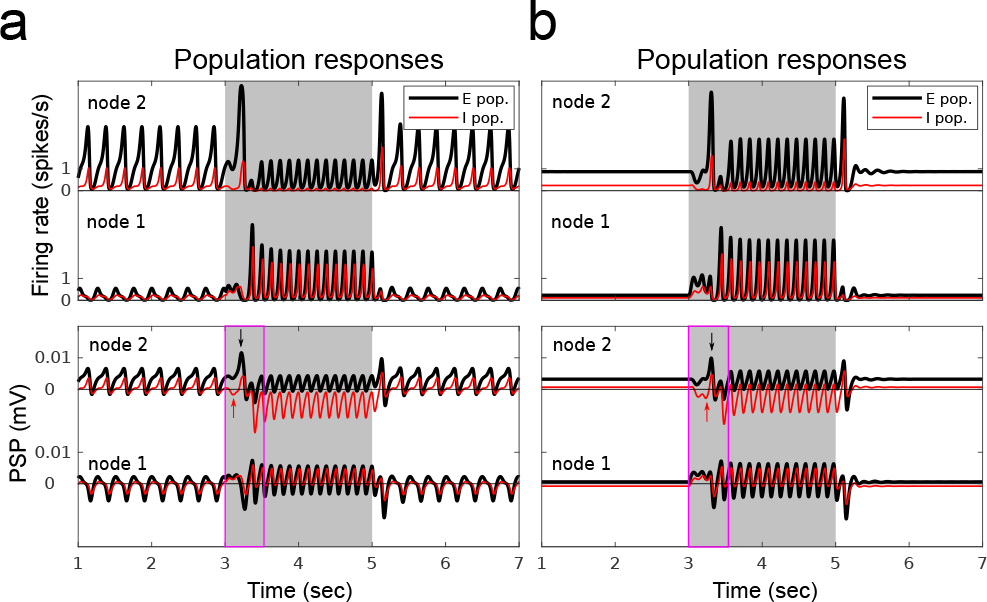
The generation of On response. **a** In the simulation settings, a prolonged stimulus of 2000 ms is fed to the R node (node 1) in a two-node network. The inter-node connections *W* is chosen from the *W* solutions that give rise to inhibited-On&Off responses in the C node (node 2) (see Figure 2b). **b** Node responses to the stimulus (gray period) are shown in firing rates (upper plot) and in PSPs (lower plot). As an inhibited-On&Off response, the time course 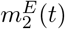 (black curve) shows lower amplitude during stimulus and transient peaks at the onset and offset of stimulus. To understand the generation of On response in population *E*_2_, the magenta rectangle indicate the period of transient disinhibition where population *I*_2_ is transiently inhibited by population *I*_1_ (red arrow), and population *E*_2_ peaks right after the transient disinhibition (black arrow). **c** and **d** show a similar example for sustained-On&Off response, where the generation of On response is also due to the transient disinhibition.

#### The generation of Off responses

Off responses proceeded by inhibited activity (i.e. the inhibited-Off responses) have been widely accepted to arise from post-inhibitory rebound that is related to the intrinsic conductance property of the neuron membranes [44]. However, the generation of the Off responses that follow sustained activity (i.e. the sustained-Off responses) cannot be simply explained by the post-inhibition mechanism (see review in [106,43]). Here we examine under which conditions the inhibited-Off and sustained-Off responses might arise at the network level.

In Figure 7, the population *E*_2_ shows Off responses for both cases: the inhibited and sustained activity during stimulus. In the simulations, both inhibited-Off and sustained-Off responses result from the same mechanism. As shown in Figure 7b and e, the Off response comes in two steps. First, the population *I*_2_ receives strong inhibition from population *I*_1_ during stimulus (reflected by the negative PSP 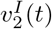 during t=3000 to 5000 ms). Second, the population *E*_2_ activity peaks before *I*_2_ recovers after stimulus offset (the transient peak 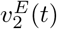 during t=5000 to 5100 ms). The occurrence of Off responses can be also represented by phase portraits shown in Figure 7c and f. The trajectories of the phase portraits show how 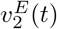 and 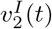 evolve interactively. When there is only background input, *E*_2_ and *I*_2_ oscillate in the normal steady state (the counter-clockwise blue trajectories) where *E*_2_ excites *I*_2_, and *I*_2_ inhibits *E*_2_. During stimulus presentation, *E*_2_ and *I*_2_ oscillate in a reversed steady state (the clockwise green trajectories) where *E*_2_ has an additional inhibitory effect on *I*_2_ through the pathway *E*_2_ → *I*_1_ → *I*_2_, and *I*_2_ has an additional disinhibitory effect on *E*_2_ through the pathway *I*_2_ → *I*_1_ → *E*_2_, due to the involvement of active *I*_1_ during stimulus. The Off responses are depicted by the magenta trajectories during the transition from the reversed steady state to the normal steady state.

**Fig. 7.**
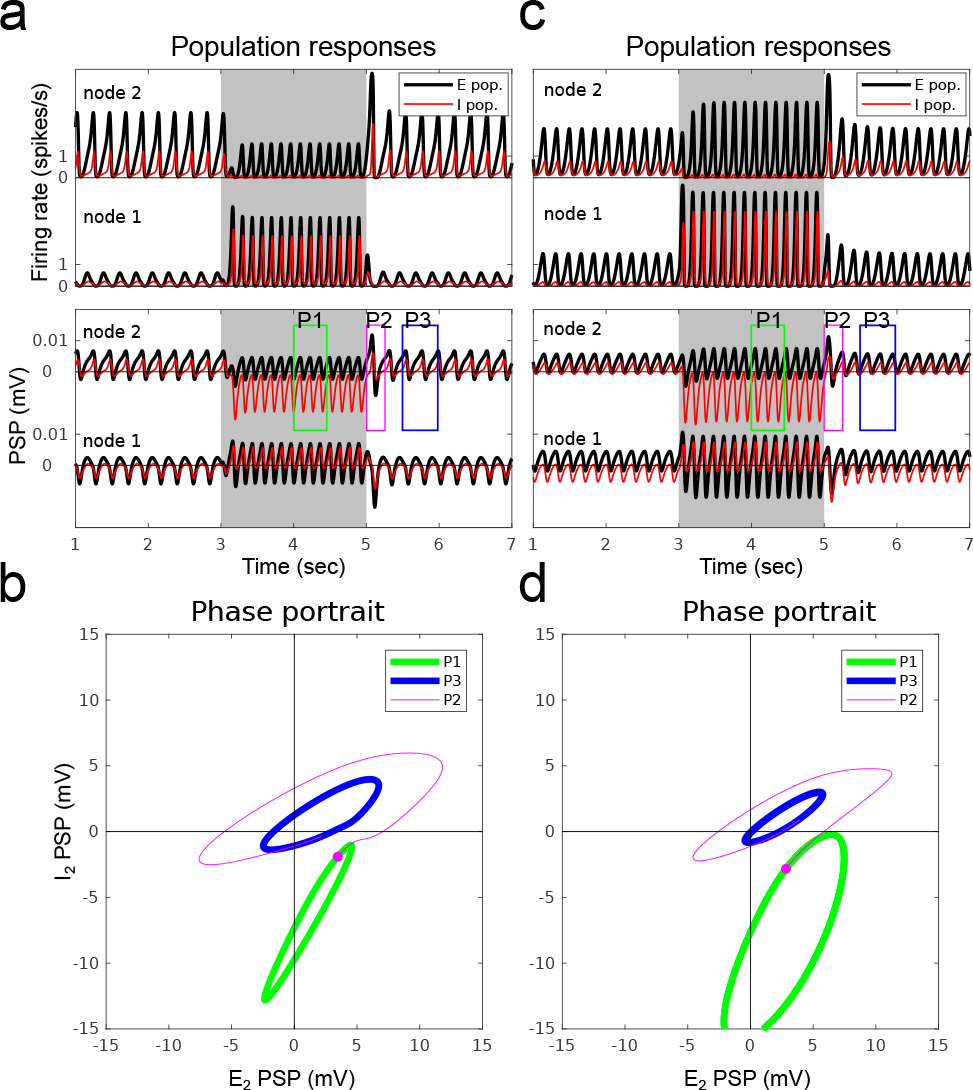
The generation of the Off response. **a** In the simulation settings, a prolonged stimulus of 2000 ms is fed to the R node (node 1) in a two-node network. The inter-node connections *W* is chosen from the *W* solutions that give rise to inhibited-Off responses in the C node (node 2) (see Figure 2b). **b** Node responses to the stimulus (the gray period) are shown in firing rates (upper plot) and in PSPs (lower plot). As an inhibited-Off response, the time course 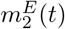 (black curve) shows lower amplitude during stimulus and a transient peak at the offset of stimulus. Population *I*_2_ is strongly inhibited during stimulus, which is reflected by the negative PSP of population 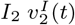 (red curve in the green rectangle). The disinhibition is followed by the Off response in population *E*_2_ thereafter (black curve in the magenta rectangle). **c** Phase portraits (*P*1: during stimulus, *P*2: offset of stimulus, *P*3: post stimulus) of node 2. The phase portrait *P*3 (i.e., when there is only background input) runs counter-clockwise, and the phase portrait *P*1 (i.e., during stimulus) shifts downward and runs clockwise, reflecting the strong inhibition on *I*_2_. The phase portrait *P*2 shows the transient trajectory of transition from *P*1 to *P*3. The magenta dot denotes the time of stimulus offset. **d** The simulation settings for a sustained-Off response. **e** The firing rate 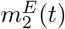 shows higher amplitude during stimulus and a transient peak at the offset of stimulus. Same as in **b**, population *I*_2_ is strongly inhibited during stimulus, which is then followed by the Off response. **f** The phase portraits are similar as in **c** except that the amplitude of 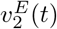 is larger during *P*1 than *P*3. The two examples show that the generation of Off responses are not relevant to the sustained or inhibited activity in *E*_2_, but the inhibition on *I*_2_ during stimulus.

The simulations provide clues for the underlying neural mechanism. The inter-node connection *W*^*II*^ is critical for a network to give rise to the Off responses because some inhibitory population *I*_2_ has to be inhibited (i.e. disinhibition) first. The inter-node connection *W*^*EI*^ is important to maintain the network in the working state (e.g., the reversed steady state) otherwise the network gets ‘overheated’ during disinhibition. With these structural prerequisites, the excitatory population *E*_2_ may show a transient Off response before the inhibitory population *I*_2_ catches up again after the stimulus offset.

The timing of the stimulus offset (i.e. the initial point in the state space when the transition begins) and other parameters that alter the trajectories of the two steady states (such as the stimulus intensity, and the settings of *W*^*EE*^ and *W*^*IE*^) also affect the generation of Off response, but these factors are not critical. Moreover, the inhibited activity during stimulus is not critical for the generation of the Off response at network level (cf., it is necessary in the post-inhibitory mechanism at cellular level). As can be seen in Figure 7f, the amplitude of 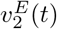 during stimulus (green trajectory) can be larger compared to no stimulus (blue trajectory).

#### Factors influencing the On/Off responses

We consider the effect of three factors with respect to the generation of On/Off responses: (1) external input to inhibitory populations, (2) blockage of NMDA receptor channels, and (3) synaptic adaptation. More specifically, we checked how these three factors influence the distribution of *W* solutions in the two-node network.

Since disinhibition plays an important role in the generation of both On and Off responses as illustrated in the above simulations (Figure 6 and 7), it is interesting to see the contribution of external input to the inhibitory population *I*_1_. In condition II, the external connection *W*^*IX*^ is set to zero in comparison with the default setting *W*^*IX*^ = 0.5*W*^*EX*^ (condition I).

The NMDA-r antagonist MK-801 is found to reduce inhibition during stimulation and thus reduce the Off responses [4]. NMDA-r antagonists are also known to reduce the amplitude of the MMN [55]. In condition III, we mimic the effect of NMDA-r antagonists by reducing the connection strength *W*^*EE*^ by 25% and reducing *W*^*IE*^ by 50%. The difference in reduction applied to the two connections is based on the fact that excitatory synapses on inhibitory neurons are mainly covered by NMDA channels and therefore are more sensitive to NMDA-r antagonists than the excitatory synapses on excitatory neurons [74]. The setting of external connections remains the same as the default setting. redNote that, in principle, both conditions II and III might be sensitive to NMDA-r antagonists. So, if NMDA-r antagonists are indeed the cause of reduced connection strengths to inhibitory populations, the effect in condition II and III in Figure 8h should occur simultaneously. Other effects by NMDA-r antagonists such as the changes in NMDA currents, synaptic plasticity and synaptic time constant are not included.

**Fig. 8.**
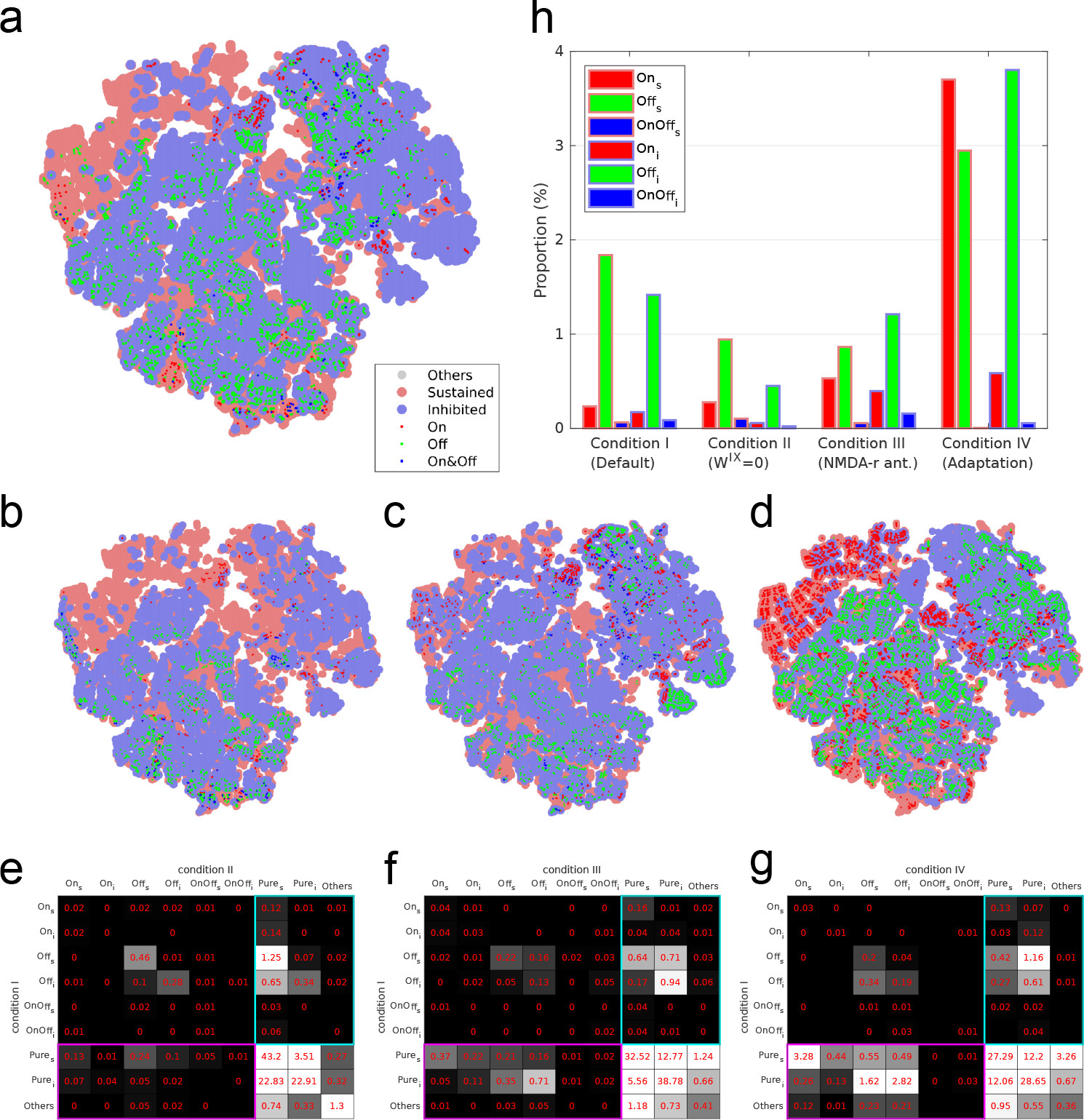
The effect of factors influencing the strength of connections W on the occurrence of On/Off responses. The *W* solutions of On/Off responses projected to 2D plane under **a** condition I: default, **b** condition II: *W*^*IX*^ = 0, **c** condition III: NMDA-r antagonist, and **d** condition IV: synaptic adaptation. Dots with different colors and sizes represent different response types. **e** The contingency table of *W* solutions for condition I vs. II. The value in each cell of the table (in red, with grayscale background) is the number of *W* solutions over total number of scanned *W*s. The subscripts *s* and *i* stand for sustained and inhibited types. A cell without a value means no *W* solution in that case. The cyan and magenta rectangles highlight the *W* solutions of On/Off types under one condition but not under the other. **f** Condition I vs. III. **g** Condition I vs. IV. **h** The bar chart represents the proportions of *W* solutions of On/Off types under the four conditions.

The phenomenon of synaptic adaptation is ubiquitous and has been suggested to be one of the mechanisms underlying deviance detection. Since we suggest that deviance-related responses can be interpreted as the change detection responses to regularity representation, it is good to know whether synaptic adaptation promotes the emergence of change detectors. To consider synaptic adaptation in the simulation, the intra- and inter-node connections *W*^*EE*^ are modulated by a synaptic efficacy term *a* as described in Equation 17. Note that as described in Equation 9, the external input via *W*^*EX*^ to the excitatory populations is not affected by synaptic adaptation.

The responses of two-node network with a range of inter-node connections *W*s (same as in Example I in Section 3.1) are simulated, and each *W* is assigned to one of the nine types of responses (i.e., eight On/Off types and type 9: others. See also Figure 2b) under four conditions: (I) the default condition, where synaptic adaptation is not applied, and *W*^*IX*^ = 0.5*W*^*EX*^, *W*^*IX*^ = 0, (III) *W*^*EE*^ = 0 is reduced by 25% and *W*^*IE*^ = 0 is reduced by 50%, and (IV) synaptic adaptation is applied. To visualize the result, the *W* solutions of types 1 to 9 are projected to 2D plane (Figure 8a-d). The number of W solutions under the four condition are summarized in the contingency table (Figure 8e-g) and the bar chart (Figure 8h).

The bar chart (Figure 8h) shows that the number of *W* solutions of Off types in condition II are reduced compared to condition I. Most of the Off types under condition I turn to Pure types under condition II (e.g., sustained-Off → sustained-Pure among 1.25% of the scanned Ws. See the cyan rectangle in Figure 8e). This suggests that the external connection *W*^*IX*^ is supportive for the generation of Off responses, because the *I*_1_-to-*I*_2_ disinhibition is enhanced due to the external input via *W*^*IX*^.

In condition III the number of *W* solutions of Off types are reduced, but the number of *W* solutions of On types are slightly increased compared to condition I (Figure 8h). This is in line with the experimental result that NMDA-r antagonist reduce the Off responses but the On responses are not affected [37, 99].

In condition IV, the number of *W* solutions of both On and Off types are greatly increased (Figure 8h). Many of the Pure types under condition I turn into On and Off types under condition III (e.g., 3.28%: sustained-Pure → sustained-On; 1.62%: inhibited-Pure → sustained-Off; 2.82%: inhibited-Pure → inhibited-Off. See the magenta rectangle in Figure 8f). This suggests that synaptic adaptation greatly promotes the emergence of change detectors. To see how synaptic adaptation alters the network responses, Figure 9a-c show three examples of altered responses due to synaptic adaptation. The three examples show typical type transitions from condition I to condition IV.

**Fig. 9.**
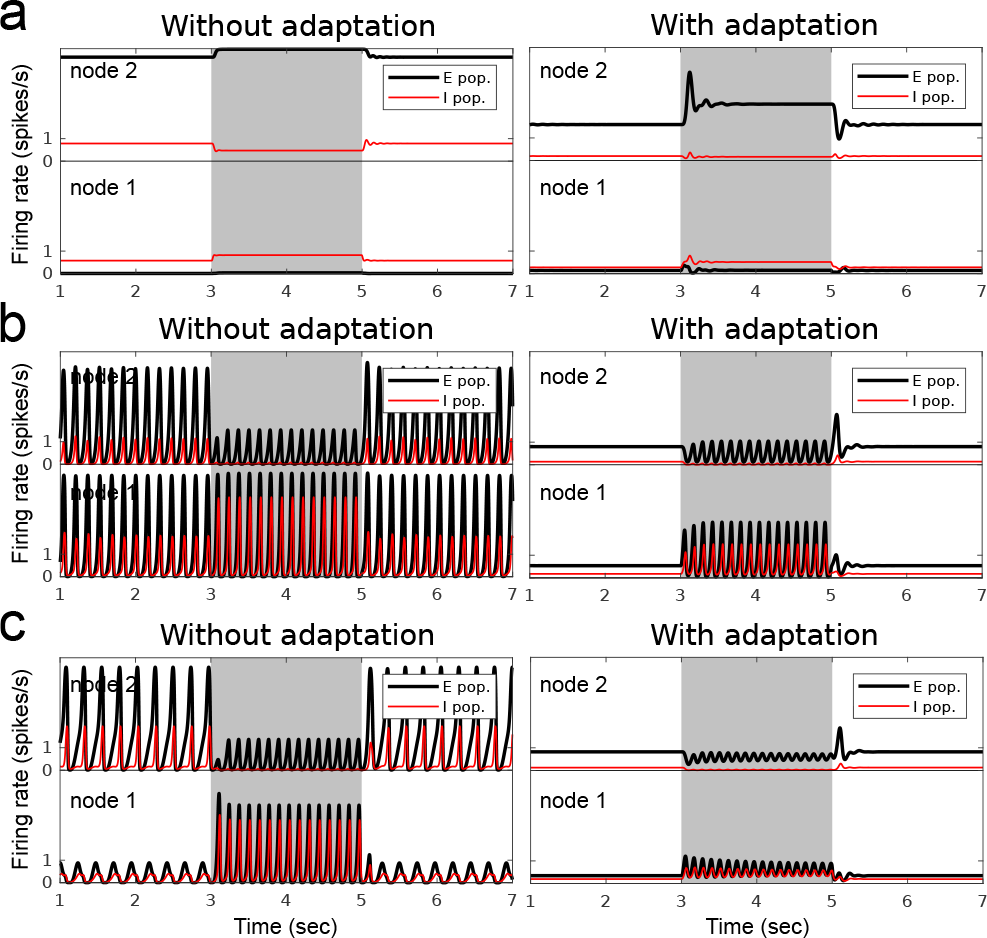
Network responses without and with synaptic adaptation. **a** Example of sustained-Pure type response in population *E*_2_ under condition I (i.e., no adaptation) turning into sustained-On type under condition IV (i.e., when synaptic adaptation is applied on *W*^*EE*^). **b** Example of inhibited-Pure type turning into sustained-Off type. **c** Example of inhibited-Pure type turning into inhibited-Off type.

## 4 Discussion

We propose a *generic deviance detection* principle based on the fact that many deviance-related responses can be observed in the cortex without clear evidence of functionally-specific wiring patterns. The mechanism suggests that the reciprocal wirings in the cortex give rise to the emergence of change detectors that respond to the abrupt change in regular features. With this notion, the deviance-related responses observed in the cortex such as the cortical On/Off responses, the cortical OSR and the MMN are regarded as responses of change detectors at different levels of abstraction.

The simulation examples demonstrate the network responses can resemble the properties of cortical On/Off responses (Figure 2 and 3), the cortical OSR (Figure 4), and the MMN (Figure 5). We then study the wiring patterns in the network that support the generation of On/Off responses (Figure 6 and 7). The results suggest that the inhibitory to inhibitory connections are important for both On and Off responses, which implies that these deviance-related responses are closely related to disinhibition. In the simulations that mimic the effect of NMDA-r antagonists and synaptic adaptation, the results show that NMDA-r antagonists suppress the Off responses and slightly promote the On responses, and synaptic adaptation generally boosts both On and Off responses (Figure 8 and 9). Below we provide our viewpoints regarding the questions raised in the introduction. Some testable predictions by our model are listed at the end of the discussion.

### Different processes in regularity formation, but same mechanism in change detection

The generic deviance detection principle suggests that the *change detection* may rely on a common neural mechanism (i.e. the local reciprocal wiring), while the *regularity formation* may, depending on the level of abstraction, require different brain resources and time to collect relevant information.

There are a number of dissimilarities among different deviance-related responses, which are, as we will discuss hereunder, mainly due to differences in the process of regularity formation. We take the differences between cortical OSR and MMN as an example. In terms of temporal window of integration (TWI), a pitch MMN can be elicited by traditional oddball paradigms even when the SOA is larger than 500 ms [77,6], while the estimated length of TWI for cortical OSR is much shorter (l60-l70 ms) [107]. In terms of attention, it is suggested that fast and slow periodic sequences elicit cortical OSRs by two different mechanisms: The fast OSR (periodicity > 5 Hz) is elicited automatically, while the slow OSR (periodicity < 2 Hz) requires the involvement of attention [41]. For example, the slow OSR can be elicited at large SOAs such as 800 ms in [25]; and 1000 to 2000 ms in [15]. The need for attention suggests that cortical OSR and MMN are different processes [60]. In terms of required repetition, a successful elicitation of MMN needs only two or three repetitions for simple feature-repetition regularities [8,18,91,105], while the cortical OSR requires up to 9 repetitions in a train for a successful elicitation [29]. The above observations suggest for different processes that are related to the degree of difficulty in regularity formation.

There are several similarities among the deviance-related responses that support the notion of a common mechanism for change detection. In terms of latency, the peak latencies of cortical On/Off responses, cortical OSR, and MMN all fall in the range of 100-200 ms [2,57,73,107,70]. In terms of spatial distribution, the sources of cortical Off response and MMN are similar. As revealed in animal studies, the sources of Off responses appear to be in the non-tonotopic area that is adjacent to the tonotopic area [92,4]. In studies of dense mapping of MMN, the pitch MMN was reported to be generated in the secondary auditory area (or spreading more widely over the core and belt areas), which is separate from the sources of P1 and N1 at the core areas (A1 and AAF) [86,67]. The cortical responses to the onset, offset, and pitch change in a continuous stimulus share similar topography and temporal profile, as suggested in the EEG/MEG studies [110,111,62]. These observations support the notion of a common neural substrate of change detection for different deviance-related responses.

The deviance-related responses also show similarities in their dependency on some factors regarding the regularities (e.g., probability of deviant, randomness in SOA, number of repetition, effect of the NMDA-r antagonists) and the deviance magnitude (e.g., the sharpness in temporal, spectral, contextual changes).

### The recurrent nature of the intracortical wiring makes the change detection ubiquitous

Functionally speaking, the ubiquity of change detection facilitates the perceptual representation in the hierarchy. The edge information at all levels provided by the local change detectors augments the representational space. Such information compression may also contribute to energy saving. In this sense, the change detectors are more like high-pass filters than comparators that subtract top-down signals from the bottom-up signals. The abundant recurrent wiring patterns in the cortex provide a suitable environment for the emergence of change detectors. We take the diversity of cortical On/Off responses [16,20,100] as an example. Even though these responses could originate from the feedforward mixture of the non-cortical On/Off responses at earlier stages such as the thalamus, midbrain and brainstem, the cortex provides more abundant chances for the emergence of On/Off responses. In example simulation I, we demonstrate that various types of On/Off responses can be generated by different inter-node connections (Figure 2). In example simulation II, we further demonstrate that for a specific connection setting, the difference in input ratios to nodes gives rise to distinct onset and offset FRFs (Figure 3). The *W* solutions of On/Off responses projected to the 2D plane (Figure 2c) also provide explanation to the diverse (and spatially clustered) cell responses observed in auditory cortex in awake mice, as shown in Figure 5 in [20]. These results suggest that change detection is a basic and ubiquitous operation in the cortex.

We then study the generation of On and Off responses. The On responses can be due to a transient disinhibition (i.e., a quick and light inhibition on the inhibitory population of the change detector) before the network reaches the steady state (Figure 6). The Off responses are always due to a release from long-lasting disinhibition (i.e., a long and strong inhibition on the inhibitory population of the change detector) before the network comes back to the steady state without stimulus (Figure 7), in line with the rebound after inhibition hypothesis [92,27]. We suggest that the inhibitory to inhibitory connections are a key ingredient for change detection.

### NMDA-r antagonists dampen the deviance-related responses

We suggest that the NMDA-r antagonists could generally dampen the deviance-related responses through three aspects: (1) voltage-dependency, (2) synaptic plasticity and (3) E/I balance. First, the NMDA-r antagonists block the voltage-dependent NMDA channels and reduce the additional NMDA currents that reflect mismatch signals [37]. Second, the NMDA-r antagonists damage the spike-timing-dependent plasticity (STDP) and hamper the ability of regularity formation [4,38,98]. Third, the NMDA-r antagonists alter the connection patterns and E/I balance. Blocking NMDA receptors leads to decreased activities in the GABAergic interneurons and increased pyramidal excitation, because the GABAergic interneurons are tenfold more sensitive to the NMDA-r antagonists than the pyramidal neurons [74,23].

The adaptation-based and prediction-based models of MMN agree on the voltage-dependency aspect and suggest that the reduced MMN amplitude is due to the reduction in NMDA currents [53,102,101]. The prediction-based models also mention the need for STDP to form prediction signals [102,101]. In addition to these two aspects, our simulation results show that the altered E/I balance as an effect of NMDA-r antagonists can reduce the emergence of change detectors. In condition III (Figure 8c), we reduce the strengths of *W*^*EE*^ and *W*^*IE*^ by 25% and 50% respectively and count again the number of each On/Off types in the scanned range of inter-node connection *W*s. The number of Off types is decreased and the number of On types is slightly increased relative to the default setting (Figure 8h). The results suggest that the NMDA-r antagonists may dampen the cortical Off response, cortical OSR and MMN.

We cannot draw further quantitative conclusion on the effect of NMDA-r antagonists because the uniform search range of Ws in the simulation is just a simplification, and the exact proportion of strength reduction due to NMDA-r antagonists is not available. The setting of 25% and 50% in connection strength reduction in condition III was arbitrary so that a single node still oscillates under a certain range of input intensity, which eliminates the case when the nodes get saturated and no On/Off responses are not generated at all. The time constant *τ*_*e*_ due to the blockage of NMDA channels was not modified in the simulation in order to focus on the effect of *W* change.

### Synaptic adaptation facilitates change detection

Synaptic adaptation is a pervasive short-term synaptic plasticity that is considered as a mechanism underlying deviance detection, in the sense that a rare stimulus triggers stronger neural activities via un-adapted pathways. Given the pervasiveness of synaptic adaptation, we are interested in how it affects the behavior of the change detector in the simulation. In the simulation of condition IV (Figure 8d), the strength of *W*^*EE*^ is modulated by short-term adaptation according to the activities of presynaptic excitatory populations. After scanning through the *W*s, we found that the number of *W* solutions of both On and Off types are increased compared with the default condition (Figure 8g). Many *W* solutions of Pure type turn into On and Off types when synaptic adaptation is applied (as examples in Figure 9). We suggests that the synaptic adaptation facilitates change detection by bringing many otherwise Pure responses (usually reflected by saturated activities in the excitatory populations) to either On or Off responses.

### The OSR is not just sustained resonance

The OSR differentiates itself from the Off response by its peak latencies that are proportional to the SOA in repetitive stimuli, which reflects the role of temporal expectancy. The models that claim to account for the OSR utilize either an *adaptive approach* [94] or *population coding approach* [52] to maintain a short continuation of neural activities (i.e., sustained resonances) that preserve the periodicity of the repetitive stimuli. However, the sustained resonances alone cannot fulfil all observations in terms of peak amplitude and peak latency of the response: (1) For the peak amplitude, the OSR cannot simply rely on the sustained resonance since the amplitude of OSR can be stronger than the evoked response during entrainment [29]; (2) For the peak latency, there should a constant delay upon one more period at the end of the stimuli [2,84], but the sustained resonance rises exactly after one more period at the end of the stimuli. Therefore, even though the sustained resonance is time-locked to the next stimulus, there seem to be extra neural circuits responsible for the extra constant delay in peak latency and the stronger peak amplitude than the evoked responses. In the example simulation III, we demonstrate that the simulated OSR solves the two issues mentioned above (Figure 4). Our model suggests that the cortical OSR can be interpreted as the cortical Off response after the end of sustained resonance. The simulation result is also in line with the finding that there is pre-activated response at the time of expected onset followed by a mismatch response [2,9,79].

### The OSR is not prediction signal

The *omission paradigm* is often used to differentiate the contribution of *adaptation* and *prediction* in MMN generation. This is based on the assumption that the OSR could not arise without any stimulus without the involvement of active prediction. Interestingly, the models based on either the adaptation or the prediction hypothesis interpret the OSR as essentially different from the MMN triggered by classic oddball paradigm. In the adaptation-based model, the OSR is regarded as the rebound response (i.e. the sustained resonance) rather than delayed N1 [52,53]. In the prediction-based model, the OSR is regarded as pure prediction signal that originates from the memory unit rather than prediction error [102]. Both interpretations imply that the OSR is essentially different from the MMN because no additional NMDA current is generated. The problem is that neither the rebound response nor the prediction signal do explain the two observations in terms of amplitude and latency mentioned above. As demonstrated in the example simulations III and IV (Figure 4 and 5), we suggest that the cortical OSR and MMN are essentially the same, both being the activities of change detectors.

The *cross-modal omission paradigm* is also used to emphasize the need for prediction. The brain can predict the upcoming event (e.g. handclap sound) from the preceding events in another modality (e.g. silent handclap video, or self-paced button press), and an OSR is triggered if the expected stimulus is omitted. In a motor-auditory (MA) paradigm, the participants show OSRs when the sound that was expected to be initiated by the self-paced button press is omitted [79]. In a visual-auditory (VA) paradigm, an OSR is elicited by occasionally omitting the sound that accompanied a handclap video [90]. So far, the cross-modal OSRs have not been considered by the existing computational models. How does the generic deviance detection principle view the OSRs in these cross-modal paradigms that seem to be bound to an active predicting process? Here we provide a different view point. First, the prediction is likely to be supported by the association between the cross-modal events (e.g., handclap video or button press, followed by a sound stimulus) that has to be paired or learned (e.g., by Hebbian learning) in advance either via direct or indirect connections. The existence of association is reflected by the pre-activation at 40 to 80 ms in the auditory cortex elicited by a visual event [90] or by a motor event [79,90]. There is no preactivation in the auditory cortex in the random condition where the button press is followed by a randomly selected sound from 48 samples in the MA paradigm and there is also no OSR thereafter [79], suggesting that there are not enough trials to associate the button press to all 48 sound samples. Second, due to the pre-activation in the auditory cortex, the MA and VA paradigms can then be regarded as a classic oddball paradigm where the standard is a ‘weak-strong’ sound pair and the deviant is a ‘weak-omission’ sound pair. In this sense, the cross-modal omission paradigm resembles an ‘intensity MMN’ or ‘duration MMN’ paradigms rather than an omission paradigm. This analogy explains why OSRs are elicited in the VA and MA conditions but not in auditory-only condition (like a classic omission paradigm) [90] because the SOAs (average 1155 ms) in the paradigm is above the temporal window of integration (TWI) for temporal feature such as periodicity but still within the TWI for identity features such as intensity and duration. The analogy can be verified if the VA and MA conditions fail to elicit ‘omission’ responses when the SOAs are larger than TWI for identify features. Based on this analogy, the deviance detection that takes place in the auditory cortex stands alone from the process of association, which explains why the pre-activation does not differ between the 50% and 12% chance of sound omission, while the mismatch response following the pre-activation depends on the proportion of omission trials for both VA and MA conditions [90], because association is less likely to be reduced by the 50% omissions, but deviance detection relies much on probability. Taken together, given the pre-activation via association and the analogy to classic MMN paradigm, the computational models that account for classic MMN (e.g., either prediction-based or not) could potentially also account for the mismatch responses in cross-modal omission paradigm. From the viewpoint of generic deviance detection principle, the process of deviance detection (including regularity formation and change detection) takes place locally in the auditory cortex, even in the case of cross-modal VA and MA paradigms.

### Testable predictions

In terms of the location of response: (1) The cortical Off response, cortical OSR, and MMN should show similar laminar profiles, for example, sink in layer 2/3 [37]. (2) Inhibited activity of inhibitory interneurons near the location of the deviance response should be observed during stimulus presentation (regularity formation). Taking the pitch MMN for an example (assuming cortical area A has best frequency (BF) of standard tone A, area B has BF of deviant tone B, and area X is the location of MMN), the inhibitory interneurons in area X should be inhibited by tone A. In addition, area X can be a broader area (which may still include area B) that surrounds area A. In terms of the effect of NMDA-r antagonists: (1) The cortical OSR should be sensitive to the NMDA-r antagonists as are the other MMNs. (2) The amplitude of entrainment to periodic stimuli in omission paradigms should be also reduced by NMDA-r antagonists. Note: this prediction may have been partially supported by impaired delta entrainment in patients with schizophrenia [49].

## Acknowledgements

We would like to thank Dr. Alejandro Tabas for the helpful suggestions.

## References

1. Amenedo, E., Escera, C.: The accuracy of sound duration representation in the human brain determines the accuracy of behavioural perception. European Journal of Neuroscience 12(7), 2570–2574 (2000)

2. Andreou, L.V., Griffiths, T.D., Chait, M.: Sensitivity to the temporal structure of rapid sound sequencesan meg study. Neuroimage 110, 194–204 (2015)

3. Auksztulewicz, R., Barascud, N., Cooray, G., Nobre, A.C., Chait, M., Friston, K.: The cumulative effects of predictability on synaptic gain in the auditory processing stream. Journal of Neuroscience 37(28), 6751–6760 (2017)

4. Baba, H., Tsukano, H., Hishida, R., Takahashi, K., Horii, A., Takahashi, S., Shibuki, K.: Auditory cortical field coding long-lasting tonal offsets in mice. Scientific reports 6, 34421 (2016)

5. Barascud, N., Pearce, M.T., Griffiths, T.D., Friston, K.J., Chait, M.: Brain responses in humans reveal ideal observer-like sensitivity to complex acoustic patterns. Proceedings of the National Academy of Sciences 113(5), E616–E625 (2016)

6. Bartha-Doering, L., Deuster, D., Giordano, V., am Zehnhoff-Dinnesen, A., Dobel, C.: A systematic review of the mismatch negativity as an index for auditory sensory memory: From basic research to clinical and developmental perspectives. Psychophysiology 52(9), 1115–1130 (2015)

7. Behrend, O., Brand, A., Kapfer, C., Grothe, B.: Auditory response properties in the superior paraolivary nucleus of the gerbil. Journal of neurophysiology 87(6), 2915–2928 (2002)

8. Bendixen, A., Roeber, U., Schröger, E.: Regularity extraction and application in dynamic auditory stimulus sequences. Journal of Cognitive Neuroscience 19(10), 1664–1677 (2007)

9. Bendixen, A., Schröger, E., Winkler, I.: I heard that coming: event-related potential evidence for stimulus-driven prediction in the auditory system. Journal of Neuroscience 29(26), 8447–8451 (2009)

10. Boh, B., Herholz, S.C., Lappe, C., Pantev, C.: Processing of complex auditory patterns in musicians and non-musicians. PLoS One 6(7), e21458 (2011)

11. Brannon, E.M., Roussel, L.W., Meck, W.H., Woldorff, M.: Timing in the baby brain. Cognitive Brain Research 21(2), 227–233 (2004)

12. Bullock, T.H., Hofmann, M.H., Nahm, F.K., New, J.G., Prechtl, J.C.: Event-related potentials in the retina and optic tectum of fish. Journal of Neurophysiology 64(3), 903–914 (1990)

13. Bullock, T.H., Karamürsel, S., Achimowicz, J.Z., McClune, M.C., Başsar-Eroglu, C.: Dynamic properties of human visual evoked and omitted stimulus potentials. Electroencephalography and clinical neurophysiology 91(1), 42–53 (1994)

14. Bullock, T.H., Karamürsel, S., Hofmann, M.H.: Interval-specific event related potentials to omitted stimuli in the electrosensory pathway in elasmobranchs: an elementary form of expectation. Journal of Comparative Physiology A 172(4), 501–510 (1993)

15. Busse, L., Woldorff, M.G.: The erp omitted stimulus response to no-stim events and its implications for fast-rate event-related fmri designs. Neuroimage 18(4), 856–864 (2003)

16. Chimoto, S., Kitama, T., Qin, L., Sakayori, S., Sato, Y.: Tonal response patterns of primary auditory cortex neurons in alert cats. Brain research 934(1), 34–42 (2002)

17. Colin, C., Hoonhorst, I., Markessis, E., Radeau, M., De Tourtchaninoff, M., Foucher, A., Collet, G., Deltenre, P.: Mismatch negativity (mmn) evoked by sound duration contrasts: an unexpected major effect of deviance direction on amplitudes. Clinical neurophysiology 120(1), 51–59 (2009)

18. Cowan, N., Winkler, I., Teder, W., Näätänen, R.: Memory prerequisites of mismatch negativity in the auditory event-related potential (erp). Journal of Experimental Psychology: Learning, Memory, and Cognition 19(4), 909 (1993)

19. Dehmel, S., Kopp-Scheinpflug, C., Dörrscheidt, G.J., Rübsamen, R.: Electrophysiological characterization of the superior paraolivary nucleus in the mongolian gerbil. Hearing research 172(1–2), 18–36 (2002)

20. Deneux, T., Kempf, A., Daret, A., Ponsot, E., Bathellier, B.: Temporal asymmetries in auditory coding and perception reflect multi-layered nonlinearities. Nature communications 7, 12682 (2016)

21. Felix, R.A., Fridberger, A., Leijon, S., Berrebi, A.S., Magnusson, A.K.: Sound rhythms are encoded by postinhibitory rebound spiking in the superior paraolivary nucleus. Journal of Neuroscience 31(35), 12566–12578 (2011)

22. Gai, Y.: On and off inhibition as mechanisms for forward masking in the inferior colliculus: a modeling study. American Journal of Physiology-Heart and Circulatory Physiology (2016)

23. Grunze, H.C., Rainnie, D.G., Hasselmo, M.E., Barkai, E., Hearn, E.F., McCarley, R.W., Greene, R.W.: Nmda-dependent modulation of ca1 local circuit inhibition. Journal of Neuroscience 16(6), 2034–2043 (1996)

24. Guo, Y., Burkard, R.: Onset and offset responses from inferior colliculus and auditory cortex to paired noise-bursts: inner hair cell loss. Hearing research 171(1–2), 158–166 (2002)

25. Halgren, E., Baudena, P., Clarke, J.M., Heit, G., Liégeois, C., Chauvel, P., Musolino, A.: Intracerebral potentials to rare target and distractor auditory and visual stimuli. i. superior temporal plane and parietal lobe. Electroencephalography and clinical neurophysiology 94(3), 191–220 (1995)

26. He, J.: Corticofugal modulation on both on andoff responses in the nonlemniscal auditory thalamus of the guinea pig. Journal of Neurophysiology 89(1), 367–381 (2003)

27. He, J., Hashikawa, T., Ojima, H., Kinouchi, Y.: Temporal integration and duration tuning in the dorsal zone of cat auditory cortex. Journal of Neuroscience 17(7), 2615–2625 (1997)

28. Herholz, S.C., Lappe, C., Pantev, C.: Looking for a pattern: an meg study on the abstract mismatch negativity in musicians and nonmusicians. BMC neuroscience 10(1), 42 (2009)

29. Horváth, J., Müller, D., Weise, A., Schröger, E.: Omission mismatch negativity builds up late. Neuroreport 21(7), 537–541 (2010)

30. Hsiao, F.J., Cheng, C.H., Liao, K.K., Lin, Y.Y.: Corticocortical phase synchrony in auditory mismatch processing. Biological Psychology 84(2), 336–345 (2010)

31. Hsu, W.Y., Cheng, C.H., Lin, H.C., Liao, K.K., Wu, Z.A., Ho, L.T., Lin, Y.Y.: Memory-based mismatch response to changes in duration of auditory stimuli: An meg study. Clinical Neurophysiology 121(10), 1744–1750 (2010)

32. Jacobsen, T., Schröger, E.: Measuring duration mismatch negativity. Clinical Neurophysiology 114(6), 1133–1143 (2003)

33. Jacobsen, T., Schröger, E., Horenkamp, T., Winkler, I.: Mismatch negativity to pitch change: varied stimulus proportions in controlling effects of neural refractoriness on human auditory event-related brain potentials. Neuroscience letters 344(2), 79–82 (2003)

34. Jansen, B.H., Rit, V.G.: Electroencephalogram and visual evoked potential generation in a mathematical model of coupled cortical columns. Biological cybernetics 73(4), 357–366 (1995)

35. Jansen, B.H., Zouridakis, G., Brandt, M.E.: A neurophysiologically-based mathematical model of flash visual evoked potentials. Biological cybernetics 68(3), 275–283 (1993)

36. Jaramillo, M., Paavilainen, P., Näätänen, R.: Mismatch negativity and behavioural discrimination in humans as a function of the magnitude of change in sound duration. Neuroscience Letters 290(2), 101–104 (2000)

37. Javitt, D.C., Steinschneider, M., Schroeder, C.E., Arezzo, J.C.: Role of cortical n-methyl-d-aspartate receptors in auditory sensory memory and mismatch negativity generation: implications for schizophrenia. Proceedings of the National Academy of Sciences 93(21), 11962–11967 (1996)

38. Javitt, D.C., Sweet, R.A.: Auditory dysfunction in schizophrenia: integrating clinical and basic features. Nature Reviews Neuroscience 16(9), 535 (2015)

39. Joachimsthaler, B., Uhlmann, M., Miller, F., Ehret, G., Kurt, S.: Quantitative analysis of neuronal response properties in primary and higher-order auditory cortical fields of awake house mice (m us musculus). European Journal of Neuroscience 39(6), 904–918 (2014)

40. Karamürsel, S., Bullock, T.H.: Dynamics of event-related potentials to trains of light and dark flashes: responses to missing and extra stimuli in elasmobranch fish. Electroencephalography and clinical neurophysiology 90(6), 461–471 (1994)

41. Karamürsel, S., Bullock, T.H.: Human auditory fast and slow omitted stimulus potentials and steady-state responses. International Journal of Neuroscience 100(1–4), 1–20 (2000)

42. Knösche, T.R., Lattner, S., Maess, B., Schauer, M., Friederici, A.D.: Early parallel processing of auditory word and voice information. NeuroImage 17(3), 1493–1503 (2002)

43. Kopp-Scheinpflug, C., Sinclair, J.L., Linden, J.F.: When sound stops: offset responses in the auditory system. Trends in neurosciences 41(10), 712–728 (2018)

44. Kopp-Scheinpflug, C., Tozer, A.J., Robinson, S.W., Tempel, B.L., Hennig, M.H., Forsythe, I.D.: The sound of silence: ionic mechanisms encoding sound termination. Neuron 71(5), 911–925 (2011)

45. Kuchenbuch, A., Paraskevopoulos, E., Herholz, S.C., Pantev, C.: Effects of musical training and event probabilities on encoding of complex tone patterns. BMC neuroscience 14(1), 51 (2013)

46. Kujala, T., Kallio, J., Tervaniemi, M., Näätänen, R.: The mismatch negativity as an index of temporal processing in audition. Clinical Neurophysiology 112(9), 1712–1719 (2001)

47. Kulesza Jr, R.J., Spirou, G.A., Berrebi, A.S.: Physiological response properties of neurons in the superior paraolivary nucleus of the rat. Journal of neurophysiology (2003)

48. Large, E.W., Almonte, F.V., Velasco, M.J.: A canonical model for gradient frequency neural networks. Physica D: Nonlinear Phenomena 239(12), 905–911 (2010)

49. Lee, M., Sehatpour, P., Hoptman, M.J., Lakatos, P., Dias, E.C., Kantrowitz, J.T., Martinez, A.M., Javitt, D.C.: Neural mechanisms of mismatch negativity dysfunction in schizophrenia. Molecular psychiatry 22(11), 1585 (2017)

50. Lehmann, A., Arias, D.J., Schönwiesner, M.: Tracing the neural basis of auditory entrainment. Neuroscience 337, 306–314 (2016)

51. Maaten, L.v.d., Hinton, G.: Visualizing data using t-sne. Journal of machine learning research 9(Nov), 2579–2605 (2008)

52. May, P., Tiitinen, H.: Human cortical processing of auditory events over time. NeuroReport 12(3), 573–577 (2001)

53. May, P.J., Tiitinen, H.: Mismatch negativity (mmn), the deviance-elicited auditory deflection, explained. Psychophysiology 47(1), 66–122 (2010)

54. May, P.J., Westö, J., Tiitinen, H.: Computational modelling suggests that temporal integration results from synaptic adaptation in auditory cortex. European Journal of Neuroscience 41(5), 615–630 (2015)

55. Näätänen, R., Kähkönen, S.: Central auditory dysfunction in schizophrenia as revealed by the mismatch negativity (mmn) and its magnetic equivalent mmnm: a review. International Journal of Neuropsychopharmacology 12(1), 125–135 (2009)

56. Näätänen, R., Lehtokoski, A., Lennes, M., Cheour, M., Huotilainen, M., Iivonen, A., Vainio, M., Alku, P., Ilmoniemi, R.J., Luuk, A., et al.: Language-specific phoneme representations revealed by electric and magnetic brain responses. Nature 385(6615), 432 (1997)

57. Näätänen, R., Paavilainen, P., Alho, K., Reinikainen, K., Sams, M.: Do event-related potentials reveal the mechanism of the auditory sensory memory in the human brain? Neuroscience letters 98(2), 217–221 (1989)

58. Näätänen, R., Paavilainen, P., Rinne, T., Alho, K.: The mismatch negativity (mmn) in basic research of central auditory processing: a review. Clinical neurophysiology 118(12), 2544–2590 (2007)

59. Näätänen, R., Syssoeva, O., Takegata, R.: Automatic time perception in the human brain for intervals ranging from milliseconds to seconds. Psychophysiology 41(4), 660–663 (2004)

60. Ng, K.K., Penney, T.B.: Probing interval timing with scalp-recorded electroencephalography (eeg). In: Neurobiology of Interval Timing, pp. 187–207. Springer (2014)

61. Nishihara, M., Inui, K., Morita, T., Kodaira, M., Mochizuki, H., Otsuru, N., Motomura, E., Ushida, T., Kakigi, R.: Echoic memory: investigation of its temporal resolution by auditory offset cortical responses. PloS one 9(8), e106553 (2014)

62. Nishihara, M., Inui, K., Motomura, E., Otsuru, N., Ushida, T., Kakigi, R.: Auditory n1 as a change-related automatic response. Neuroscience research 71(2), 145–148 (2011)

63. Novitski, N., Huotilainen, M., Tervaniemi, M., Näätänen, R., Fellman, V.: Neonatal frequency discrimination in 250–4000-hz range: Electrophysiological evidence. Clinical Neurophysiology 118(2), 412–419 (2007)

64. Novitski, N., Tervaniemi, M., Huotilainen, M., Näätänen, R.: Frequency discrimination at different frequency levels as indexed by electrophysiological and behavioral measures. Cognitive Brain Research 20(1), 26–36 (2004)

65. Paavilainen, P.: The mismatch-negativity (mmn) component of the auditory event-related potential to violations of abstract regularities: a review. International journal of psychophysiology 88(2), 109–123 (2013)

66. Phillips, D.P., Hall, S., Boehnke, S.: Central auditory onset responses, and temporal asymmetries in auditory perception. Hearing research 167(1–2), 192–205 (2002)

67. Pincze, Z., Lakatos, P., Rajkai, C., Ulbert, I., Karmos, G.: Separation of mismatch negativity and the n1 wave in the auditory cortex of the cat: a topographic study. Clinical Neurophysiology 112(5), 778–784 (2001)

68. Prechtl, J.C., Bullock, T.H.: Event-related potentials to omitted visual stimuli in a reptile. Electroencephalography and Clinical Neurophysiology 91(1), 54–66 (1994)

69. Qin, L., Chimoto, S., Sakai, M., Wang, J., Sato, Y.: Comparison between offset and onset responses of primary auditory cortex on-off neurons in awake cats. Journal of neurophysiology (2007)

70. Raij, T., McEvoy, L., Mäkelä, J.P., Hari, R.: Human auditory cortex is activated by omissions of auditory stimuli. Brain research 745(1–2), 134–143 (1997)

71. Ramón, F., Hernández, O.H., Bullock, T.H.: Eventrelated potentials in an invertebrate: crayfish emit omitted stimulus potentials. Journal of experimental biology 204(24), 4291–4300 (2001)

72. Recanzone, G.H.: Response profiles of auditory cortical neurons to tones and noise in behaving macaque monkeys. Hearing research 150(1–2), 104–118 (2000)

73. Rinne, T., Särkkä, A., Degerman, A., Schröger, E., Alho, K.: Two separate mechanisms underlie auditory change detection and involuntary control of attention. Brain research 1077(1), 135–143 (2006)

74. Rujescu, D., Bender, A., Keck, M., Hartmann, A.M., Ohl, F., Raeder, H., Giegling, I., Genius, J., McCarley, R.W., Möller, H.J., et al.: A pharmacological model for psychosis based on n-methyl-d-aspartate receptor hypofunction: molecular, cellular, functional and behavioral abnormalities. Biological psychiatry 59(8), 721–729 (2006)

75. Ruusuvirta, T., Lipponen, A., Pellinen, E., Penttonen, M., Astikainen, P.: Auditory cortical and hippocampal-system mismatch responses to duration deviants in urethane-anesthetized rats. PloS one 8(1), e54624 (2013)

76. Saha, D., Sun, W., Li, C., Nizampatnam, S., Padovano, W., Chen, Z., Chen, A., Altan, E., Lo, R., Barbour, D.L., et al.: Engaging and disengaging recurrent inhibition coincides with sensing and unsensing of a sensory stimulus. Nature communications 8, 15413 (2017)

77. Sams, M., Hari, R., Rif, J., Knuutila, J.: The human auditory sensory memory trace persists about 10 sec: neuromagnetic evidence. Journal of cognitive neuroscience 5(3), 363–370 (1993)

78. Sams, M., Paavilainen, P., Alho, K., Näätänen, R.: Auditory frequency discrimination and event-related potentials. Electroencephalography and Clinical Neurophysiology/Evoked Potentials Section 62(6), 437–448 (1985)

79. SanMiguel, I., Saupe, K., Schröger, E.: I know what is missing here: electrophysiological prediction error signals elicited by omissions of predicted what but not when. Frontiers in human neuroscience 7, 407 (2013)

80. Scholl, B., Gao, X., Wehr, M.: Nonoverlapping sets of synapses drive on responses and off responses in auditory cortex. Neuron 65(3), 412–421 (2010)

81. Schönwiesner, M., Novitski, N., Pakarinen, S., Carlson, S., Tervaniemi, M., Naatanen, R.: Heschl’s gyrus, posterior superior temporal gyrus, and mid-ventrolateral prefrontal cortex have different roles in the detection of acoustic changes. Journal of Neurophysiology (2007)

82. Schröger, E., Paavilainen, P., Näätänen, R.: Mismatch negativity to changes in a continuous tone with regularly varying frequencies. Electroencephalography and Clinical Neurophysiology/Evoked Potentials Section 92(2), 140–147 (1994)

83. Schwartz, G., Harris, R., Shrom, D., Berry II, M.J.: Detection and prediction of periodic patterns by the retina. Nature neuroscience 10(5), 552 (2007)

84. Schwartz, G.W., Berry II, M.J.: Sophisticated temporal pattern recognition in retinal ganglion cells. Journal of neurophysiology (2008)

85. Shiga, T., Althen, H., Cornella, M., Zarnowiec, K., Yabe, H., Escera, C.: Deviance-related responses along the auditory hierarchy: Combined ffr, mlr and mmn evidence. PloS one 10(9), e0136794 (2015)

86. Shiramatsu, T.I., Kanzaki, R., Takahashi, H.: Cortical mapping of mismatch negativity with deviance detection property in rat. PLoS One 8(12), e82663 (2013)

87. da Silva, F.L.: Functional localization of brain sources using eeg and/or meg data: volume conductor and source models. Magnetic resonance imaging 22(10), 1533–1538 (2004)

88. Spiegler, A., Kiebel, S.J., Atay, F.M., Knösche, T.R.: Bifurcation analysis of neural mass models: Impact of extrinsic inputs and dendritic time constants. NeuroImage 52(3), 1041–1058 (2010)

89. Spiegler, A., Knösche, T.R., Schwab, K., Haueisen, J., Atay, F.M.: Modeling brain resonance phenomena using a neural mass model. PLoS computational biology 7(12), e1002298 (2011)

90. Stekelenburg, J.J., Vroomen, J.: Predictive coding of visual–auditory and motor-auditory events: An electrophysiological study. Brain research 1626, 88–96 (2015)

91. Sussman, E.S., Horváth, J., Winkler, I., Orr, M.: The role of attention in the formation of auditory streams. Perception & psychophysics 69(1), 136–152 (2007)

92. Takahashi, H., Nakao, M., Kaga, K.: Cortical mapping of auditory-evoked offset responses in rats. Neuroreport 15(10), 1565–1569 (2004)

93. Tervaniemi, M., Schröger, E., Saher, M., Näätänen, R.: Effects of spectral complexity and sound duration on automatic complex-sound pitch processing in humans– a mismatch negativity study. Neuroscience Letters 290(1), 66–70 (2000)

94. Thivierge, J.P., Cisek, P.: Spiking neurons that keep the rhythm. Journal of computational neuroscience 30(3), 589–605 (2011)

95. Tiitinen, H., May, P., Reinikainen, K., Näätänen, R.: Attentive novelty detection in humans is governed by pre-attentive sensory memory. Nature 372(6501), 90 (1994)

96. Toufan, R., Moossavi, A., Aghamolaei, M., Ashayeri, H.: Topographic comparison of mmn to simple versus pattern regularity violations: The effect of timing. Neuroscience research 112, 20–25 (2016)

97. Tse, C.Y., Penney, T.B.: Preattentive timing of empty intervals is from marker offset to onset. Psychophysiology 43(2), 172–179 (2006)

98. Uhlhaas, P.J., Singer, W.: Abnormal neural oscillations and synchrony in schizophrenia. Nature reviews neuroscience 11(2), 100 (2010)

99. Umbricht, D., Schmid, L., Koller, R., Vollenweider, F.X., Hell, D., Javitt, D.C.: Ketamine-induced deficits in auditory and visual context-dependent processing in healthy volunteers: implications for models of cognitive deficits in schizophrenia. Archives of general psychiatry 57(12), 1139–1147 (2000)

100. Volkov, I., Galazjuk, A.: Formation of spike response to sound tones in cat auditory cortex neurons: interaction of excitatory and inhibitory effects. Neuroscience 43(2–3), 307–321 (1991)

101. Wacongne, C.: A predictive coding account of mmn reduction in schizophrenia. Biological psychology 116, 68–74 (2016)

102. Wacongne, C., Changeux, J.P., Dehaene, S.: A neuronal model of predictive coding accounting for the mismatch negativity. Journal of Neuroscience 32(11), 3665–3678 (2012)

103. van Wassenhove, V., Lecoutre, L.: Duration estimation entails predicting when. Neuroimage 106, 272–283 (2015)

104. Werner, B., Cook, P.B., Passaglia, C.L.: Complex temporal response patterns with a simple retinal circuit. Journal of neurophysiology (2008)

105. Winkler, I., Karmos, G., Naätänen, R.: Adaptive modeling of the unattended acoustic environment reflected in the mismatch negativity event-related potential. Brain research 742(1–2), 239–252 (1996)

106. Xu, N., Fu, Z.Y., Chen, Q.C.: The function of offset neurons in auditory information processing. Translational Neuroscience 5(4), 275–285 (2014)

107. Yabe, H., Tervaniemi, M., Sinkkonen, J., Huotilainen, M., Ilmoniemi, R.J., Naätänen, R.: Temporal window of integration of auditory information in the human brain. Psychophysiology 35(5), 615–619 (1998)

108. Yago, E., Corral, M.J., Escera, C.: Activation of brain mechanisms of attention switching as a function of auditory frequency change. Neuroreport 12(18), 4093–4097 (2001)

109. Yago, E., Escera, C., Alho, K., Giard, M.H.: Cerebral mechanisms underlying orienting of attention towards auditory frequency changes. Neuroreport 12(11), 2583–2587 (2001)

110. Yamashiro, K., Inui, K., Otsuru, N., Kakigi, R.: Change-related responses in the human auditory cortex: An meg study. Psychophysiology 48(1), 23–30 (2011)

111. Yamashiro, K., Inui, K., Otsuru, N., Kida, T., Kakigi, R.: Automatic auditory off-response in humans: an meg study. European Journal of Neuroscience 30(1), 125–131 (2009)

112. Yaron, A., Hershenhoren, I., Nelken, I.: Sensitivity to complex statistical regularities in rat auditory cortex. Neuron 76(3), 603–615 (2012)

